# Ablation of SYK kinase from primary human Natural Killer cells via CRISPR/Cas9 enhances cytotoxicity and cytokine production

**DOI:** 10.1101/2022.05.29.493889

**Authors:** James D. Dahlvang, Jenna K. Dick, Jules A. Sangala, Emily J. Pomeroy, Kristin M. Snyder, Juliette M. Moushon, Claire E. Thefaine, Jianming Wu, Sara E. Hamilton, Martin Felices, Jeffrey S. Miller, Bruce Walcheck, Beau R. Webber, Branden S. Moriarity, Geoffrey T. Hart

## Abstract

Cytomegalovirus (CMV) infection alters natural killer (NK) cell phenotype and function toward a more memory-like immune state. These cells, termed adaptive NK cells, typically express CD57 and NKG2C but lack expression of the Fc receptor γ chain (Gene: *FCER1G*, FcRγ), PLZF, and SYK. Functionally, adaptive NK cells display enhanced antibody-dependent cellular cytotoxicity (ADCC) and cytokine production. However, the mechanism behind this enhanced function is unknown. To understand what drives cytotoxicity and cytokine production in adaptive NK cells, we optimized a CRISPR/Cas9 system to ablate genes from primary human NK cells. ADCC by human NK cells is exclusively mediated by the CD16A (*FcγRIIIA*) signaling apparatus, which includes FcRγ, CD3ζ, SYK, SHP-1, ZAP-70, and the transcription factor PLZF. We ablated the genes encoding these molecules and tested subsequent ADCC and cytokine production. We found that ablating the FcRγ chain caused a modest increase in TNFα production. Ablation of PLZF did not enhance ADCC or cytokine production. Importantly, SYK kinase ablation significantly enhanced both cytotoxicity and cytokine production, while ZAP-70 kinase ablation diminished function. Ablation of the phosphatase SHP-1 resulted in mixed effects on function, with NK cells demonstrating enhanced cytotoxicity but reduced cytokine production. These results indicate that the enhanced cytotoxicity and cytokine production of CMV-induced adaptive NK cells is more likely due to the loss of SYK than the lack of FcRγ or PLZF. The lack of SYK expression may limit SHP-1-mediated inhibition of CD16A signaling, leading to enhanced cytotoxicity and cytokine production. In addition to providing mechanistic answers about CMV-induced adaptive NK cell functionality, our results indicate that NK chimeric antigen receptor (CAR) therapeutics that invoke ADCC signaling molecules (e.g., CD3ζ chain) may benefit from ablating SYK, while maintaining ZAP-70, to increase functionality.

## Introduction

Natural killer (NK) cells are innate lymphocytes that can kill both cancerous and infected cells (Abel et al., 2018; Lanier, 2008). In peripheral blood, NK cells comprise 5 to 20% of lymphocytes, mainly surveying the bloodstream for potential target cells (Burrack et al., 2019; Mujal et al., 2021; Perera Molligoda Arachchige, 2021). NK cells kill target cells and signal neighboring cells through a variety of mechanisms, including natural cytotoxicity (e.g., missing self), FasL- or TRAIL-induced apoptosis, antibody-dependent cellular cytotoxicity (ADCC), and cytokine production. During ADCC and cytokine production, NK cells first recognize the Fc region of IgG bound to a target cell with CD16A (*FcγRIIIA*). CD16A utilizes signaling adapter molecules including the Fc receptor γ chain (FcRγ) and/or the CD3 ζ chain (CD3ζ) for signal transduction (Lanier et al., 1991). Despite their seemingly redundant functions, FcRγ and CD3ζ have distinct features. For example, FcRγ contains one immunoreceptor tyrosine-based activation motif (ITAM), whereas CD3ζ contains three ITAMs (Love and Hayes, 2010). Crosslinking of CD16A by antibodies bound to a target cell recruits Src-family kinases, such as LCK, to phosphorylate ITAMs on FcRγ and/or CD3ζ. This phosphorylation recruits the tyrosine kinases SYK and/or ZAP-70 to ITAMs for subsequent signal transduction. SYK preferentially binds FcRγ, whereas ZAP-70 preferentially binds CD3ζ (Shiue et al., 1995). SYK and ZAP-70 then phosphorylate their respective downstream targets (e.g., PI3K, VAV1, and phospholipase Cγ isoforms) to further propagate the CD16A signaling pathway, ultimately cumulating in cytotoxicity and cytokine production (Freedman et al., 2015; Watzl and Long, 2010). While the molecules participating in CD16A-mediated signaling pathway are defined, the specific contributions of these signaling proteins to primary human NK cell function were unclear.

Circulating NK cells are not a homogenous population. Various factors, such as infection, can drive NK cell phenotypic and functional diversity (Sun et al., 2009; Yang et al., 2019). Cytomegalovirus (CMV) is the epitome of a pathogen that alters both NK cell phenotype and function. In mice, murine CMV (mCMV) infection results in expansion of a subset of NK cells expressing Ly49H, a receptor that binds mCMV peptide on MHC. This NK population persists, specifically expands upon mCMV reactivation, and has enhanced effector function, displaying adaptive-like qualities (Sun *et al*., 2009). In humans, CMV infection also results in expansion of NK cells expressing NKG2C and CD57 (Guma et al., 2004; Lopez-Verges et al., 2011; Monsivais-Urenda et al., 2010). These cells proliferate after recognizing CMV *in vitro* (Guma et al., 2006) and upon CMV reactivation in humans (Foley et al., 2012). After CMV reactivation, CD57+/NKG2C+/NKG2A- NK cells also exhibit enhanced cytokine production (Foley *et al*., 2012). These results led to the classification of CD57+/NKG2C+/NKG2A- NK cells as adaptive NK cells due to their adaptive qualities, such as longevity and antigen-specific expansion (albeit through a limited germline-encoded receptor). Adaptive NK cells are also hypothesized to play a protective role in cancer and transplant patients as a higher frequency of adaptive NK cells correlates with better clinical outcomes in individuals with reactivated CMV (Cichocki et al., 2019).

In addition to phenotypic differences, adaptive NK cells also show increases in ADCC and cytokine production compared to conventional NK cells (Hwang et al., 2012; Zhang et al., 2013). Zhang et al. showed that CMV infection was associated with loss of the FcRγ chain. FcRγ-negative (FcRγ-) NK cells showed increased degranulation (CD107a), IFNγ, and TNFα production in response to opsonized infected cells over conventional NK cells (Zhang *et al*., 2013). In addition to FcRγ deficiency, adaptive NK cells have been shown to lack both SYK kinase and PLZF (Lee et al., 2015; Schlums et al., 2015). PLZF is a transcription factor that binds the promoter of genes encoding FcRγ and SYK and likely plays a role in regulation of these genes (Correia et al., 2018; Schlums *et al*., 2015). In addition to CMV, other infectious diseases, such as HIV and malaria, can induce adaptive NK cell formation. HIV- and malaria-induced adaptive NK cells also show enhanced ADCC and cytokine production. However, these cells have a different phenotype than their CMV-induced counterparts, such as maintained SYK expression, highlighting adaptive NK cell heterogeneity (Arora et al., 2018; Hart et al., 2019; Zhou et al., 2015).

The absence of FcRγ, PLZF, and/or SYK is hypothesized to be the mechanism responsible for enhancing the ADCC function of CMV-induced adaptive NK cells. Yet it was unclear whether these molecules are responsible for enhanced function or only correlate with it. Here, we investigated the role of molecules associated with CMV-induced adaptive NK cells— including FcRγ, PLZF, and SYK—in the ADCC signaling apparatus. We tested for redundancy in the ADCC signaling pathway by targeting molecules with similar functions as FcRγ and Syk, such as CD3ζ and ZAP-70, respectively. We also sought to disentangle which molecules are associated with cytotoxic activity versus cytokine production after CD16A activation. Using a murine model could be one approach. However, murine NK cells do not perform ADCC well and the murine CD3ζ chain does not associate well with murine CD16 as an adapter protein. This limits the utility of murine signaling protein knockouts (Aguilar et al., 2022). For humans, until recently, it has not been possible to ablate genes in primary human NK cells. Therefore, to accomplish these goals, we developed a CRISPR/Cas9 gene-editing protocol for thawed human primary NK cells, ablated genes that encoded molecules in the CD16A pathway, and tested the effect of gene ablation on ADCC and cytokine production.

We show that SYK kinase ablation enhances both ADCC and cytokine production, while ZAP-70 ablation shows the opposite phenotype. SYK ablation enhanced killing more so than degranulation and the increased killing capacity was maintained for four weeks post-expansion. These results point toward a mechanism involving SYK for the enhanced function of CMV-induced adaptive NK cells. Overall, this study provides a mechanistic understanding of the enhanced ADCC function observed in CMV-induced adaptive NK cells and provides a new strategy for improving NK cell therapies by leveraging the ADCC pathway.

## Methods

### Resource Availability

#### Lead Contact

Requests for information or resources should be directed to the lead contact, Geoffrey Hart.

#### Materials Availability

This study did not generate unique reagents.

#### Data and code availability

R 4.1 (R, 2021) was used to generate Figure S1G. The code was adapted from https://biocorecrg.github.io/CRG_RIntroduction/volcano-plots.html (Bonnin, 2020) and is available at https://github.com/jamesdahlvang/volcano. All raw data is available upon request.

### Experimental Model and Subject Details

#### Primary Cell Purification and Freezing

Primary red blood cells (RBCs) and peripheral blood mononuclear cells (PBMCs) were obtained and purified from deidentified adult blood donors (Memorial Blood Center). The work done in this study was approved by the Institutional Review Board (IRB) of the University of Minnesota. The cytomegalovirus (CMV) status of each donor was reported by Memorial Blood Center. RBCs were purified from whole blood by filtration (Memorial Blood Center, 4C4300) and resuspended at 50% hematocrit in RPMI-1640 + 25 mM HEPES, L-Glutamine, and 50 mg/L Hypo-Xanthine (Kd Medical, 50-101-8907). PBMCs were isolated using Ficoll (MP Biomedicals, ICN50494) centrifugation and the remaining RBCs were lysed using ACK (Lonza, BP10-548E). Once isolated, PBMCs were counted and resuspended at 2×10^8^ cells/mL in 10mL of PBS + 2% FBS (PEAK Serum, PS-FB1) + 1 mM EDTA (SC Buffer). NK cells were then isolated from these PBMCs using a magnetic negative selection kit (STEMCELL Technologies, 17955). Briefly, PBMCs were stained with 500μl of the manufacturer’s antibody cocktail, which is less concentrated than the manufacturer’s protocol, and 30 μl of additional biotinylated anti-CD3 (STEMCELL Technologies, 18051) was added to the sample to improve purity. Samples were incubated at RT for 10 minutes. After incubation, 1 mL of magnetic beads was added to the sample, which was then vortexed 2-3 times and incubated for 10 minutes at RT. Post-incubation, 35 mL of SC Buffer was added to the cells, which were then placed on a magnet and incubated at RT for 10 minutes. After incubation, the liquid—which contained purified NK cells—was removed from the tube without touching the magnet. NK cells were then added to a new 50 mL conical and placed back on the magnet for 10 minutes at RT. After 10 minutes, purified NK cells were removed and counted. For cryopreservation, NK cells were resuspended in RMPI-1640 (Fisher, SH3002701) + 50% FBS (PEAK Serum, PS-FB1) at 10^7^ cells/mL. 500 μL of the suspension was transferred to a cryovial. When all tubes were ready to freeze, 500 μl of FBS (PEAK Serum, PS-FB1) +15% DMSO was added to cells in cryovials. The cells were immediately transferred to a pre-cooled Mr. Frosty (Thermo-Fisher, 5100-0001) and then placed in a -80° C freezer. After 24-48 hours, the cells were transferred to a vapor-phase liquid nitrogen freezer for later use.

#### K562-artificial Antigen Presenting Cell (aAPC) Cell Line Maintenance

Modified K562-aAPC cells were graciously provided by Dr. Branden Moriarity’s lab. These cells express 4-1BB ligand and IL-21 to aid NK cell proliferation (Pomeroy et al., 2020). K562-aAPCs were maintained between 1×10^5^ to 1.5×10^6^ cells/mL in RPMI-1640 (Fisher, SH3002701) + 10% FBS (PEAK Serum, PS-FB1) (RP10) + 1 μg/mL gentamicin (Sigma-Aldrich, G1272) and incubated at 37 °C with 5% CO_2_. Unless listed otherwise, all 37 °C incubations are conducted at 5% CO_2_.

#### SKOV-3 Cell Line Maintenance

Both conventional SKOV-3 and fluorescent SKOV-3 Nuclight Red cells (SKOV-3 NLR) were graciously provided by Dr. Jeffrey Miller’s lab. Once the cells became 80-95% confluent, the medium was aspirated from the flask and the cells were washed with PBS. After washing the cells, 5 mL of pre-warmed 0.05% Trypsin-EDTA (Gibco, 25300062) was added to the flask. The cells were then incubated at 37 °C for 5 minutes. Following incubation, the flask was tapped to dislodge cells from the plastic. The cells were collected and washed with 10 mL of pre-warmed McCoy’s 5A Medium (Gibco, 166600082) + 10% FBS (PEAK Serum, PS-FB1) + 1% Pen/Strep (Gibco, 1514022) (SKOV-3 Media). The resulting suspension was centrifuged, resuspended in pre-warmed SKOV-3 Media, and counted. Half of the cell suspension was used for counting. The SKOV-3 cells were then diluted to 6.5×10^4^ cells/mL with SKOV-3 media, transferred to a new flask, and incubated at 37 °C.

#### NK-92 Cell Line Maintenance

NK-92 cells expressing GFP (parent line, NK-92-GFP), CD16A and GFP (NK-92-GFP-CD16A), and CD64/16A and GFP (NK-92-GFP-CD64/16A) were graciously provided by Dr. Bruce Walcheck’s lab. NK-92-GFP-CD16A cells stably express GFP and CD16A (Binyamin et al., 2008), whereas NK-92-GFP-CD64/16A cells stably express GFP and the cytoplasmic tail of CD16A with the extracellular domain of CD64 (Snyder et al., 2018). Every two days, NK-92 cells were resuspended and centrifuged. After centrifugation, the supernatant from the cell culture was saved. The cells were then resuspended with the same volume of pre-warmed MEM-alpha (Gibco, 12561) + 10% HI-FBS (Gibco, 16140071) + 10% HI-Horse Serum (Gibco, 26050088) + 0.8% Pen/Strep (Gibco, 1514022) + 200 IU/mL IL-2 (NCI) + 0.08 mM 2-Mercaptoethanol (Gibco, 21985023) (NK-92 Media) and counted. After counting, cells were resuspended at 2×10^5^ cells/mL in 50% NK-92 media and 50% of the previous culture’s supernatant, then incubated at 37 °C. Cells were maintained between 2×10^5^ and 1.5×10^6^ cells/mL.

#### NK Cell Expansion

Cryopreserved vials of primary NK cells were thawed by placing them in a water bath at 37 °C until the vial was 95% thawed. The cells were then transferred to a 15 mL conical. Pre-warmed RP10 was added dropwise at a rate of 1 mL/minute while the conical was swirled. After adding 3 mL of RP10, 6 more mL of pre-warmed RP10 was added and the cells were centrifuged at 650 x g. The cells were washed once with RP10, centrifuged, and resuspended in 5 mL of X-VIVO^TM^ 15 (Lonza, BE02-060F) + 10% FBS (PEAK Serum, PS-FB1) (X-Vivo 10) + 2 ng/mL IL-15 (NCI). Cells were then transferred to a 6-well plate and incubated overnight. After incubation, 3×10^6^ NK cells were mixed with 6×10^6^ irradiated K562-aAPC cells in 30 mL X-Vivo 10 + 50 IU/mL IL-2 (NCI). K562-aAPCs were irradiated at 100 grays (RS-2000 X-Ray irradiator). The cells were added to a G-Rex^©^ 6-well plate (Wilson Wolf, 80240M) and incubated at 37 °C for six days.

### Methods Details

#### CRISPR/Cas9 Gene-editing

Single-guide RNAs (sgRNA) were designed using Synthego’s CRISPR Design tool (https://design.synthego.com/#/). sgRNA sequences are listed in Supplementary Table 1. Ribonucleoprotein complexes (RNPs) were made by combining 100 pmol of sgRNA (IDT) with 30 pmol of Cas9 (IDT, 1081059). If the volume of the RNP complex was less than 5 μL/sample, nuclease-free H_2_O was added after combining the RNP complex to bring the volume to 5 μL/sample. The RNP complexes were incubated at RT for 15 minutes. If multiple genes were ablated simultaneously, individual RNP complexes were made and combined after incubating for 15 minutes.

NK cells were centrifuged and resuspended in P3 Primary Cell Nucleofector^©^ Solution (Lonza, V4XP-3032) at 3.33×10^8^ cells/mL. 5×10^6^ cells were mixed with RNPs and transferred without forming bubbles to a 16-well Nucleocuvette^©^ Strip (Lonza, V4XP-3032). Unless otherwise stated, the cells were nucleofected with an Amaxa 4D-Nucleofector^TM^ (Lonza) using code CA137. Post-nucleofection, the cells were rested for 15 minutes at RT. 80 μL of pre-warmed X-Vivo 10 was added to each nucleofection well. The cells were rested for 30 more minutes at RT, then the total volume was transferred to a G-Rex^©^ 24-well plate (Wilson Wolf, 80192M) with 1.9 mL of pre-warmed X-Vivo 10. Each well was then washed with 100 μl of media, which was transferred to the cell suspension. The cells were rested once more for 30 minutes at RT before 4 mL of X-Vivo 10 + 4.5 ng/mL of IL-15 (NCI) was added to each well (3 ng/mL IL-15 final). The cells were then incubated at 37 °C for six days.

#### Degranulation and Cytokine Production Assay with Red Blood Cells as NK Targets

RBCs were opsonized with polyclonal anti-RBC rabbit antibody (Rockland, 1094139) and combined with NK cells at a ratio of 1 NK:5 RBC in RP10 + 1 μg/mL brefeldin A (BFA) + 2 µM monensin + 1:200 anti-CD107a, which measures degranulation (Biolegend, 328625). As a negative control, NK cells were also incubated with non-opsonized RBCs. The cells were then incubated at 37 °C for 5 hours. After five hours, the cells were centrifuged, washed, and stained for flow cytometry.

#### Flow Cytometry

Cells were resuspended in a master mix made of PBS, viability dye (Tonbo, 13-0865-T500), and surface stain antibodies, then incubated at RT in the dark for 20 minutes. The cells were washed and incubated in 2% formaldehyde (Thermo, 28908) at 37 °C for 10 minutes. After incubation, the cells were washed and resuspended in 0.04% Triton (Fisher, BP151-500) at RT in the dark for exactly 7 minutes. After 7 minutes, the cells were washed in PBS + 2% FBS (PEAK Serum, PS-FB1) + 2 mM EDTA (Fisher, BP2482-1) + 2% BSA (MP Biomedicals, 810681) (FACS buffer) and stained with an internal master mix in FACS buffer. Depending on the antibodies used, the cells were then incubated at RT in the dark for 30-120 minutes. After incubation, the cells were washed with FACS buffer, washed again with PBS, and resuspended in PBS for flow cytometry. Samples were run on a CytoFLEX flow cytometer (Beckman-Coulter) and analyzed via FlowJo 10.8.0 (FlowJo) to determine the extent of gene ablation and the effect of gene ablation on cell function.

#### ICE Analysis

Synthego’s Inference of CRISPR Edits (ICE) (https://ice.synthego.com/) was used to analyze gene ablation. For ICE analysis, DNA from gene-ablated and control samples was isolated using QIAGEN’s DNAeasy kit (Qiagen, 69506) according to the manufacturer’s instructions. PCR primers were designed to amplify a region of 500 base pairs (bp) around the cut site. The forward primer was designed 100-200 bp upstream from the cut site to capture it with high-quality sequencing. Additionally, we designed sequencing primers that were nested 5-50 base pairs inside the amplicon to improve sequencing efficiency. After amplifying the sgRNA cut site, approximately 100 ng of the PCR product was submitted to the UMN Genomics center along with 6.4 pmol of a sequencing primer for Sanger sequencing. The primers used are listed in Supplementary Table 1.

#### 350 Surface Protein Analysis (LEGENDScreen^TM^)

To assess the expression of approximately 350 surface proteins at once, cells were analyzed using Biolegend’s LEGENDScreen^TM^ Human PE Kit (Biolegend, 700007). Subjects were barcoded with combinations of CD45 antibodies conjugated to three fluorophores, allowing us to test 7 subjects in one well (A, B, C, AB, AC, BC, ABC). After barcoding and pooling, samples were stained for CD56, CD3, live/dead, and CD14, then acquired on a BD Fortessa flow cytometer according to manufacturer instructions.

#### DELFIA Killing Assay with SKOV-3 cells as NK Targets

DELFIA killing assays were performed according to manufacturer instructions (PerkinElmer, AD0116). Briefly, SKOV-3 (HER-2 +) cells were resuspended at 10^6^ cells/mL in SKOV-3 Media. To label the cells, 1.5 μL/mL of Bis(acetoxymethyl)-2-2:6,2 terpyridine 6,6 dicarboxylate (BATDA) was added to the suspension. The cell suspension was then incubated for 30 minutes at 37 °C. After labeling the cells, they were washed and transferred to a round-bottom 96-well plate at a concentration of 8×10^3^ cells/well. Anti-HER2 (Trastuzumab) was then added to the well at a concentration of 3 or 10 µg/mL along with either primary NK or NK-92 (effector) cells at various E:T ratios. The plate was spun at 400 x g for one minute and incubated at 37 °C for 2 hours with 5% CO_2_. After 2 hours, the plate was centrifuged at 500 x g for five minutes. 20 μL of supernatant was then transferred to a 96-well DELFIA Yellow Plate and combined with 200 μl of europium. Signal was measured by time-resolved fluorescence using a BioTek Synergy 2. To measure maximum release, BADTA-labeled target cells were incubated with 10μL of lysis buffer per the manufacturer’s instruction. To measure spontaneous lysis, BADTA-labeled SKOV-3 cells were cultured in parallel *without* effector cells or antibodies. When possible, samples were run in technical triplicates.

#### Incucyte Killing Assay with SKOV-3 cells as NK Targets

On the day prior to the killing assay, SKOV-3 NLR cells (HER-2 +) were resuspended at 4×10^4^ cells/mL in RP10. 100 µl of the cell suspension was added to each well of an Incucyte-compatible flat-bottom 96-well plate. The SKOV-3 NLR cells were then incubated overnight at 37 °C to allow them to adhere to the bottom of the well.

The following day, NK cells were resuspended at 1.2×10^6^ cells/mL in RP10. 60 µl of NK cells were added to a new 96-well plate. Anti-HER2 was then diluted to 20 µg/mL in RP10 and 60 µL of the solution was added to the NK cells. To prevent cells from localizing to the edge of the well, both the 96-well plate containing the NK cells/anti-HER2 and the plate containing the SKOV-3 NLR cells were incubated at 37 °C for 15 minutes. 100 µl of NK cells/anti-HER2 was then added to the plate containing SKOV-3 NLR cells for an effector: target (E: T) ratio around 15:1 and a final anti-HER2 concentration of 5 μg/mL. The cell mixture was then loaded into an Incucyte SX5 (Sartorius). Immediately after loading the plate, a t = 0-hour image was taken to ensure cells were not localized to the edge of the well. After confirming cells were dispersed throughout the well, four consecutive images were taken per well once per hour for 24 hours. As controls, SKOV-3 NLRs and NK cells without anti-HER2 were imaged as well as SKOV-3 NLRs without NK cells. Samples were run in technical triplicates.

### Quantification and Statistical Analysis

#### Ablation Level Cutoffs

Data points from samples with less than 25% protein ablation were excluded from all analyses to ensure the gene-ablated population was substantial enough to analyze. For expansion experiments, the cutoff was only applied at day 6 (e.g., if a sample was above 25% protein ablation at day 6 but below 25% at day 13, the data from day 13 was included). Samples between 25-49% ablation were kept for ADCC assay analysis as the gene-ablated cells can be gated on via flow cytometry. Unlike flow cytometry-based ADCC assays, killing assays utilized a target cell count readout, precluding specific analysis of gene-ablated cells. Thus, killing assay data from samples with less than 49% protein ablation were excluded.

#### LEGENDScreen^TM^ Cutoffs

Proteins that were expressed in fewer than 10% of cells from each subject at each timepoint were classified as not expressed in NK cells. These proteins were excluded from the final analysis.

#### Fold-change Data

Data were transformed using fold-change and log_2_ fold-change to ease the visualization of trends. Fold-change was calculated by dividing the gene-ablated value by the no RNP value for the measurement in question.

#### Gene Ablation Analysis

Flow cytometry was used to assess the extent of CRISPR/Cas9 gene-ablation at the protein level. To calculate the protein ablation percentage for single-gene ablations, the following formula was applied:

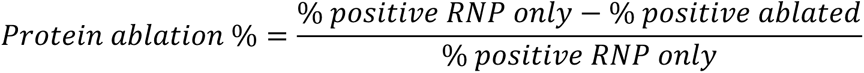

To calculate the protein ablation percentage for double-gene ablations, the following formula was used:

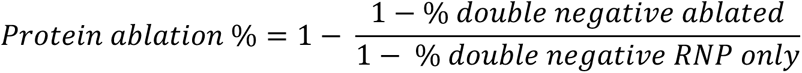

#### Incucyte Killing Assay Analysis

Incucyte Base Analysis Software (Satorius, v2019B) was used to analyze killing assays. Images with SKOV-3 NLRs + NK cells, SKOV-3 NLRs alone, and NK cells alone were chosen from the 0-hour, 12-hour, and 24-hour images. Image parameters were then adjusted to ensure the software detected SKOV-3 NLRs but not NK cells. Graphs were created by normalizing each well’s SKOV-3 NLR count at each timepoint to its time 0-hour count.

#### DELIFA Spontaneous Lysis Calculation

For each sample, specific lysis was used to report ADCC activity. The calculation for specific lysis is shown below:

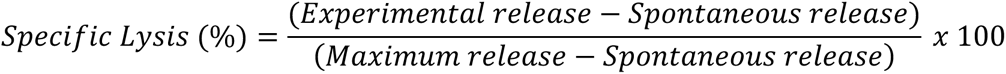

#### Graphing and Statistical Analysis

Excel 16.54 (Microsoft) was used for data transformation and statistical analysis. Schematics were created with BioRender.com. R 4.1 (R, 2021) was used to generate Figure S1G. Prism 9.1.2 (GraphPad) was used to generate figures and run statistical analyses. Illustrator 25.4.1 (Adobe) was used to draft figures for publication. Sample sizes, independent experiments run, statistical tests, and significance thresholds are listed in each figure legend.

## Results

### Development of an effective CRISPR/Cas9 gene-editing protocol to ablate genes in primary human NK cells

Previous reports of CRISPR/Cas9 in primary human NK cells have reported various platforms and settings (Huang et al., 2021; Liu et al., 2020; Naeimi Kararoudi et al., 2018; Pomeroy *et al*., 2020; Rautela et al., 2018). We aimed to optimize a CRISPR/Cas9 protocol for the Amaxa 4D system as it can be scaled up for large numbers of cells, potentially allowing for future clinical applications. To optimize our CRISPR/Cas9 protocol, we first used 24 Amaxa 4D nucleofection codes (format: XX###) to ablate FcRγ *(FCER1G)* from primary NK cells. Six days post-CRISPR, we measured gene ablation efficiency (Figure S1A) and cell recovery (Figure S1B) by flow cytometry. From these 24 electroporation codes, we picked six that balanced protein ablation and cell recovery. We also tested ‘CA137’, as it had been used for successful gene ablation in NK-92 cells (Hintz et al., 2021). We observed >70% ablation of FcRγ (Figure S1C) for each code. However, viable cell recovery varied. Of the codes tested, CA137 showed the highest average recovery at 3.3×10^6^ cells (Figure S1D). We concluded that using CA137 results in the best balance of effective gene ablation and cell recovery, leading us to use CA137 for the rest of the experiments. By evaluating a compilation of experiments that used CA137, we found that cell counts six days post-CRISPR often exceeded our starting cell count, indicating that the cells recovered well from nucleofection (Figure S1E). We also tested CRISPR/Cas9 deletion efficiency by molecular and protein-level methods to evaluate concordance. Indel analysis of sequencing data (Inference of CRISPR Edits (ICE)) and flow cytometry analysis of our target gene protein expression typically showed similar levels of gene ablation (Figure S1F). Representative gating schemes and protein ablation frequency for all data are shown in Figure S2.

We then assessed the phenotype and function of primary human NK cells post-CRISPR. To expand primary human NK cells, we incubated them for seven days by co-incubating them with IL-2, IL-15, and irradiated K562 artificial APC cells, which express 4-1BB ligand and IL-21. However, expanding human primary NK cells *in vitro* can affect NK cell phenotype (Huang *et al*., 2021). We were concerned that activation and expansion could trigger ADAM17 activity, which cleaves CD16A from the NK cell surface (Romee et al., 2013; Wang et al., 2013). Thus, to evaluate the effect of our expansion protocol on cell phenotype, we barcoded primary human NK cells from 7 subjects and examined the expression of 350 surface proteins pre- and post-expansion using Biolegend’s LEGENDScreen^TM^. As expected, we observed that many proteins were significantly up- and down-regulated during expansion (Figure S1G). However, we did not see loss of CD16A expression, which indicated that our expanded cells would retain ADCC function. We confirmed this by measuring ADCC-induced degranulation and cytokine production with an anti-RBC ADCC assay, in which red blood cells (RBCs) coated with anti-human RBC polyclonal antibodies were used as target cells. On day 6 post-CRISPR, NK cells yielded high levels of degranulation (CD107a+) and IFNγ production (Figure S1H-I). Overall, these results show that our expansion and CRISPR/Cas9 protocol generates gene-ablated primary NK cells that can be utilized in functional assays (Figure 1A).

**Figure 1.**
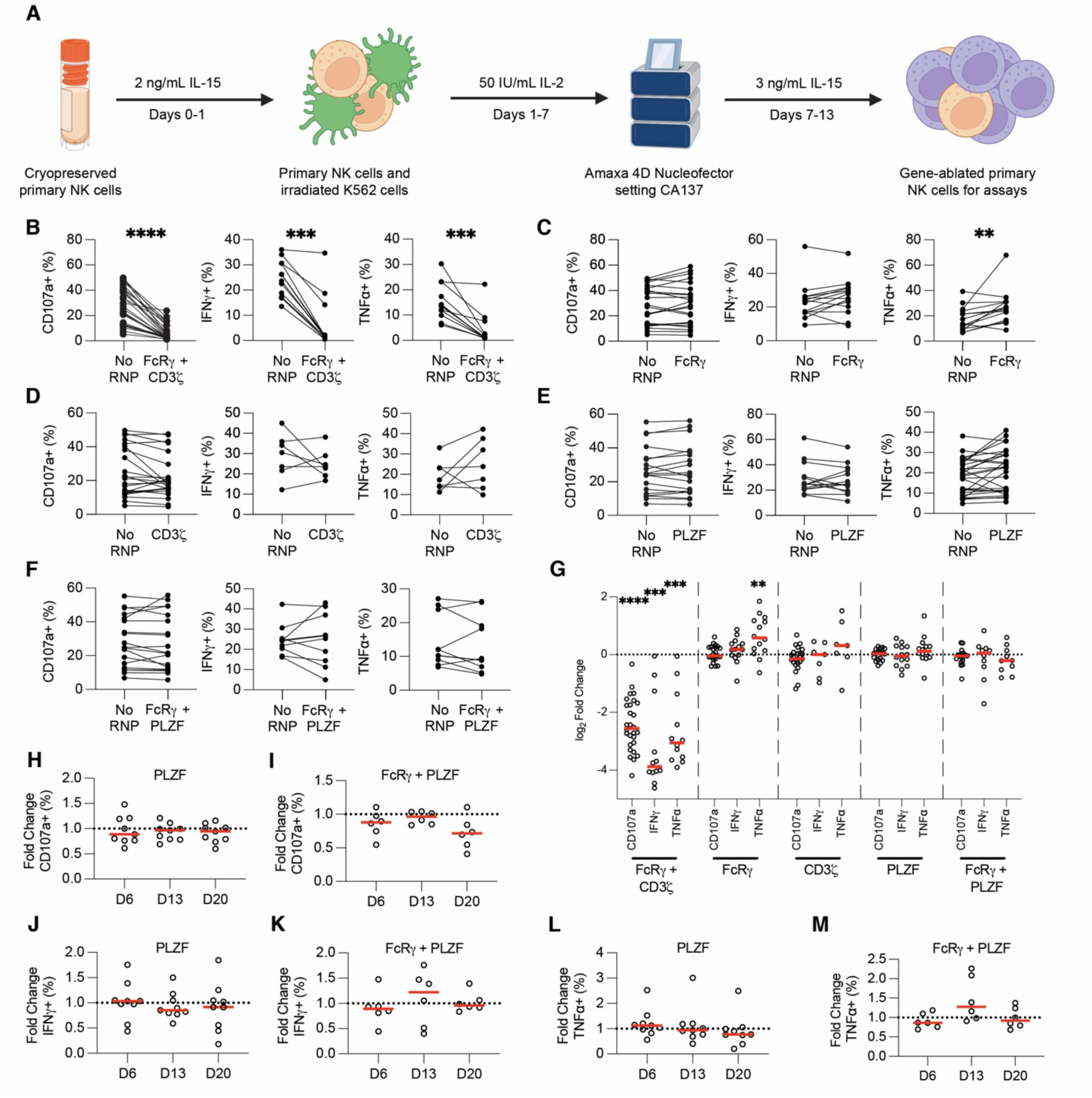
FcRγ and PLZF ablation does not affect ADCC function. Schematic depiction of CRISPR/Cas9 protocol developed and used for this study (A). Anti-RBC degranulation assays were run six days post-CRISPR/Cas9 with FcRγ/CD3ζ - (B), FcRγ- (C), CD3ζ - (D), PLZF- (E), and FcRγ-/PLZF- (F) ablated primary NK cells. CD107a, IFNγ, and TNFα production from gene-ablated and control primary NK cells was measured by flow cytometry. Each dot is one subject donor. Lines connect samples from the same donor that were nucleofected without an RNP complex (control) or with an RNP complex. Values for No RNP groups are frequencies from all NK cells, whereas values for the gene-ablated sample are only from cells negative for the protein ablated. Data from 13 (CD107a) or six (IFNγ and TNFα) independent experiments. CD107a, IFNγ, and TNFα values for each subject shown in B-F were divided by the respective no RNP sample’s value and log_2_-transformed (G). CD107a (H-I), IFNγ (J-K), and TNFα (L-M) expression was measured in PLZF- (H, J, L) and FcRγ/PLZF- (I, K, M) ablated cells at days 6, 13, and 20 post-CRISPR/Cas9. The frequency of cells positive for CD107a, IFNγ, and TNFα in the gene-ablated group was divided by the frequency of cells positive for the functional marker in the subject-matched No RNP group. Dots represent subjects and red lines represents the median. Data from three independent experiments. Wilcoxon matched-pairs signed rank test. Significance shown in G carried over from tests run for B-F. **: p < 0.01; ***: p < 0.005; ****: p < 0.001; lack of stars indicates non-significance.

### Ablating FcRγ increases TNFα, but PLZF ablation does not affect cytotoxicity or cytokine production in cellular ADCC assays

We used our CRISPR/Cas9 gene-editing protocol to test whether the loss of proteins associated with adaptive NK cells affected ADCC and cytokine production. A representative gating scheme used for analyzing ADCC assays with gene-ablated NK cells is shown in Figure S3. As a control, we first ablated both FcRγ and CD3ζ (*CD247*) from primary NK cells and then tested the gene-ablated cells in anti-RBC ADCC assays. NK cells require expression of at least one of these adapter proteins for downstream CD16A signaling, so we predicted that ablation of both genes would preclude degranulation and cytokine production. As expected, we found that simultaneous ablation of FcRγ and CD3ζ decreased CD107a, IFNγ, and TNFα expression in every subject, with an average decrease of 80-90% for each marker (Figure 1B, G). We were also interested in ablating genes from the NK-92 cell line to validate our results (Hintz *et al*., 2021; Snyder *et al*., 2018). NK-92 cells lack the FcRγ chain (Hintz *et al*., 2021), meaning ablation of CD3ζ should preclude any functional activity. Thus, we ablated CD3ζ from two NK-92 cell lines: one expressing CD16A (NK-92-GFP-CD16A) and the other expressing a recombinant high-affinity Fc receptor consisting of the CD16A cytoplasmic tail and CD64 extracellular domain (NK-92-GFP-CD65/16A). We then performed DELFIA killing assays with CD3ζ ablated NK cells and SKOV-3 cells coated with trastuzumab (anti-HER2) as targets. Ablation of CD3ζ abrogated target cell killing in both NK-92-GFP-CD16A (Figure S4A, E-H) and NK-92-GFP-CD64/16A (Figure S4C, I-L) cell lines. These controls show that our CRISPR/Cas9 sgRNAs were on-target and that differences in biological function could be revealed in both primary NK and NK-92 cells.

We next used CRISPR/Cas9 to test if adaptive NK cells have enhanced function because they lack the FcRγ chain. FcRγ only has one ITAM whereas CD3ζ contains three ITAMs. Thus, we hypothesized that if FcRγ is absent from NK cells, they will signal exclusively through CD3ζ, resulting in a stronger ADCC signal due to the increased ITAM number on CD3ζ. We also hypothesized that if the lack of the FcRγ chain enhances ADCC and cytokine production, lack of the CD3ζ chain may have the opposite effect. We ablated FcRγ and CD3ζ independently, then used the gene-ablated cells in anti-RBC ADCC assays. Surprisingly, we found that ablation of either the FcRγ or CD3ζ chain does not significantly affect degranulation or IFNγ production (Figure 1C-D, 1G). TNFα was expressed in a significantly higher frequency of FcRγ-negative cells (Figure 1C) but this was not true for CD3ζ-negative cells (Figure 1D). Overall, though ablating the FcRγ chain increases TNFα production during ADCC, neither ablation of the FcRγ nor the CD3ζ chain consistently alters ADCC function.

Adaptive NK cells also lack PLZF (*ZBTB16)* expression (Schlums *et al*., 2015). PLZF controls expression of genes associated with adaptive NK cell activity, such as FcRγ and SYK *(SYK*), leading to some speculation that PLZF may be the regulator of adaptive NK cell development (Schlums *et al*., 2015). Without PLZF, the FcRγ and other genes may not be transcribed, leading to stronger ADCC and cytokine production. We hypothesized that ablation of PLZF may trigger the adaptive NK cell transcriptional program, resulting in enhanced ADCC and cytokine production in PLZF-ablated primary NK cells. To test this hypothesis, we ablated PLZF individually or PLZF together with FcRγ (FcRγ/PLZF). We found that there was no significant difference in cytotoxicity or cytokine production with PLZF or FcRγ/PLZF-ablated cells six days post-CRISPR/Cas9 in the anti-RBC ADCC assay (Figure 1E-G). However, we were concerned that testing PLZF-ablated cells six days post-CRISPR/Cas9 may be too soon to observe an effect. Since PLZF is a transcription factor, we speculated that changes in transcription could take longer than six days post-CRISPR/Cas9 to manifest. We therefore tested PLZF and FcRγ/PLZF-ablated cells for ADCC function 6, 13, and 20 days post-CRISPR/Cas9. Neither PLZF nor FcRγ/PLZF-ablated NK cells showed enhanced degranulation, (Figure 1H-I), IFNγ (Figure 1J-K), or TNFα (Figure 1L-M) at day 13 or 20 post-CRISPR. Overall, these results showed that ablation of PLZF did not significantly affect NK cell ADCC and cytokine production in this anti-RBC ADCC assay.

### Ablations of SYK and ZAP-70 have opposite effects on degranulation and cytokine production

In addition to lacking FcRγ and PLZF, adaptive NK cells frequently lack the tyrosine kinase SYK (Lee *et al*., 2015; Schlums *et al*., 2015). Both SYK and the related ZAP-70 transduce signals from FcRγ and CD3ζ for ADCC. However, loss of SYK has been correlated with enhanced ADCC function in primary NK cells (Lee *et al*., 2015). We next tested if lack of SYK or ZAP-70 would affect ADCC and cytokine production. We ablated SYK or ZAP-70 (*ZAP70*) from primary NK cells and then examined the function of the gene-ablated cells in anti-RBC ADCC assays. We found that SYK ablation significantly enhanced degranulation, with an average increase of 7% over no RNP controls (Figure 2A, D). Of the 35 samples tested, 26 showed enhanced degranulation with SYK ablation (Figure 2A, D). In contrast, ZAP-70 ablation significantly diminished NK cell degranulation, with an average decrease of 11.9% versus no RNP controls and decreases in 15 of 22 subjects tested (Figure 2A, D). For both SYK and ZAP-70-ablation, cytokine production followed the same pattern as degranulation. SYK-ablated cells produced significantly more IFNγ (Figure 2B, D) and TNFα (Figure 2C, D) than controls, whereas ZAP-70 ablated cells expressed significantly less IFNγ (Figure 2B, D) and TNFα (Figure 2C, D).

**Figure 2.**
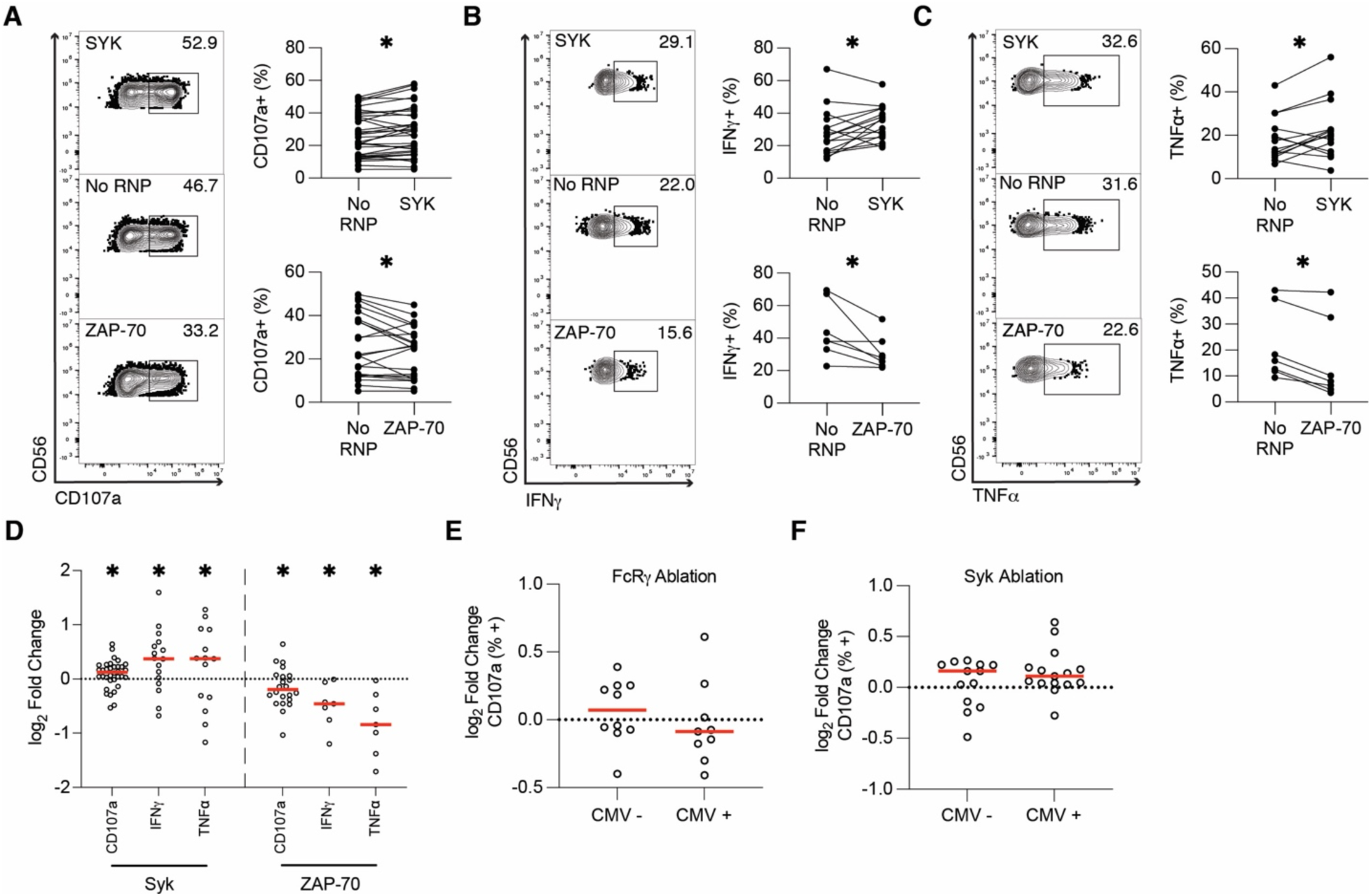
SYK ablation enhances ADCC function whereas ZAP-70 ablation hinders it. Six days post-CRISPR/Cas9, anti-RBC ADCC assays were run with SYK and ZAP-70-ablated primary NK cells. CD107a (A), IFNγ (B), and TNFα (C) production from gene-ablated and control primary NK cells was measured by flow cytometry. Flow plots show representative samples for SYK-ablated (top), No RNP (middle), and ZAP-70-ablated (bottom) primary NK cells. Each dot is one subject/donor. Lines connect samples from one subject that were nucleofected without an RNP complex (control) or with an RNP complex. Values for no RNP groups are frequencies from all NK cells, whereas values for the gene-ablated sample are only from cells negative for the protein ablated. Data from 13 (CD107a) or 7 (IFNγ and TNFα) independent experiments. CD107a, IFNγ, and TNFα values for each subject shown in A-C were divided by the respective no RNP sample’s value and log-transformed (D). Data from SYK-ablated samples in Figure 2D and FcRγ-ablated samples in Figure 1G was stratified by CMV status (E-F). Wilcoxon matched-pairs signed rank test (A-C). Significance shown in D carried over from tests run for A-C. Unpaired t-test (E-F). *: p < 0.05; lack of stars indicates non-significance.

Though CRISPR/Cas9 gene-editing with primary human NK cells is a valuable model for understanding NK cell function *in vivo*, high levels of inter-subject variability can cloud biological differences. We speculated that CMV status may account for some of this variability between donors, so we analyzed ADCC data by stratifying the tested donors based on CMV status. No significant differences were observed in ADCC between CMV-positive and CMV-negative subjects after FcRγ or SYK ablation (Figures 2E, F), indicating that CMV status did not affect the outcomes of gene ablation. Finally, to confirm the effects of SYK ablation on NK cell function, we ablated SYK in NK-92-GFP-CD16A and NK-92-GFP-CD64/16A cells and tested subsequent cytotoxicity using DELFIA killing assays with SKOV-3 as NK target cells. The results mirrored our primary NK cell ADCC results, with SYK ablation enhancing killing activity in both cell lines (Figure S4B, D-L). Overall, these results indicate that SYK ablation enhances ADCC and cytokine production, whereas ZAP-70 ablation has the opposite effect.

### Neither SYK nor ZAP-70 ablation affects surface expression of CD16A on primary NK cells

We followed these experiments by exploring the mechanism behind our results. First, we tested whether SYK or ZAP-70 ablation was altering CD16A surface expression. Stimulation can downregulate surface CD16A expression (Goodier et al., 2016; Romee *et al*., 2013; Wang *et al*., 2013). Since CD16A is the receptor that binds the IgG Fc region, up or downregulation of CD16A could result in the enhanced or diminished function we observed in gene-ablated cells. We measured surface CD16A expression in FcRγ/CD3ζ-, SYK-, and ZAP-70-ablated primary NK cells prior to ADCC. FcRγ/CD3ζ ablation diminished CD16A expression (Figure S5). This was expected, as NK cells require either FcRγ or CD3ζ to express CD16A on the cell surface (Hibbs et al., 1989; Kurosaki and Ravetch, 1989; Lanier et al., 1989). In contrast to FcRγ/CD3ζ ablation, the majority of SYK- and ZAP-70-ablated cells expressed CD16A at similar frequencies, indicating that altered ADCC function in SYK and ZAP-70-ablated primary NK cells was likely not due to changes in CD16A surface expression (Figure S5).

### SHP-1 ablation increases degranulation but decreases cytokine production

We next explored other molecules in the CD16A signaling pathway, including SHP-1. SHP-1 is a phosphatase that can dephosphorylate ZAP-70 (Brockdorff et al., 1999; Plas et al., 1996) and induce ITAM-mediated inhibition (ITAMi) (Aloulou et al., 2012; Kanamaru et al., 2008; Pinheiro da Silva et al., 2007). ITAMi is a process in which weakly ligated receptors recruit SHP-1 instead of SYK or ZAP-70 to partially phosphorylated ITAMs, allowing SHP-1 to dephosphorylate the ITAM and terminate signaling (Abram and Lowell, 2007; Ivashkiv, 2011). Interestingly, ITAMi can also be facilitated by SYK dissociating from the partially phosphorylated ITAM of a weakly occupied receptor, allowing SHP-1 to dephosphorylate the ITAM (Ben Mkaddem et al., 2014; Mkaddem et al., 2017). We hypothesized that SYK ablation may be enhancing ADCC in primary NK cells by allowing ZAP-70 to bind to FcRγ at a higher rate, thus reducing SHP-1 binding to FcRγ and subsequent ITAMi. Additionally, we hypothesized that ZAP-70 ablation may be diminishing ADCC by facilitating SYK/SHP-1 binding to FcRγ, thus facilitating ITAMi. To test this, we ablated SHP-1 (*PTPN6*) from primary NK cells and tested the SHP-1-ablated cells’ ADCC function and cytokine production. In line with our hypothesis, SHP-1 ablation robustly enhanced degranulation in the anti-RBC ADCC assay. On average, 43.55% of SHP-1-ablated samples were CD107a+, compared to 34.43% of subject-matched, no RNP samples in the assay (Figure 3A, 3D). However, both IFNγ (Figure 3B, 3D) and TNFα (Figure 3C-D) were expressed at significantly lower frequencies in SHP-1 ablated samples relative to no RNP controls. Since cytokine production reflected degranulation in both SYK and ZAP-70-ablated cells, we were surprised that SHP-1 ablation resulted in enhanced ADCC but reduced cytokine production. Overall, while our CD107a data suggest that SHP-1 could be a driving force for ITAMi, the effect of SHP-1 ablation on cytokine production is both confounding and interesting.

**Figure 3.**
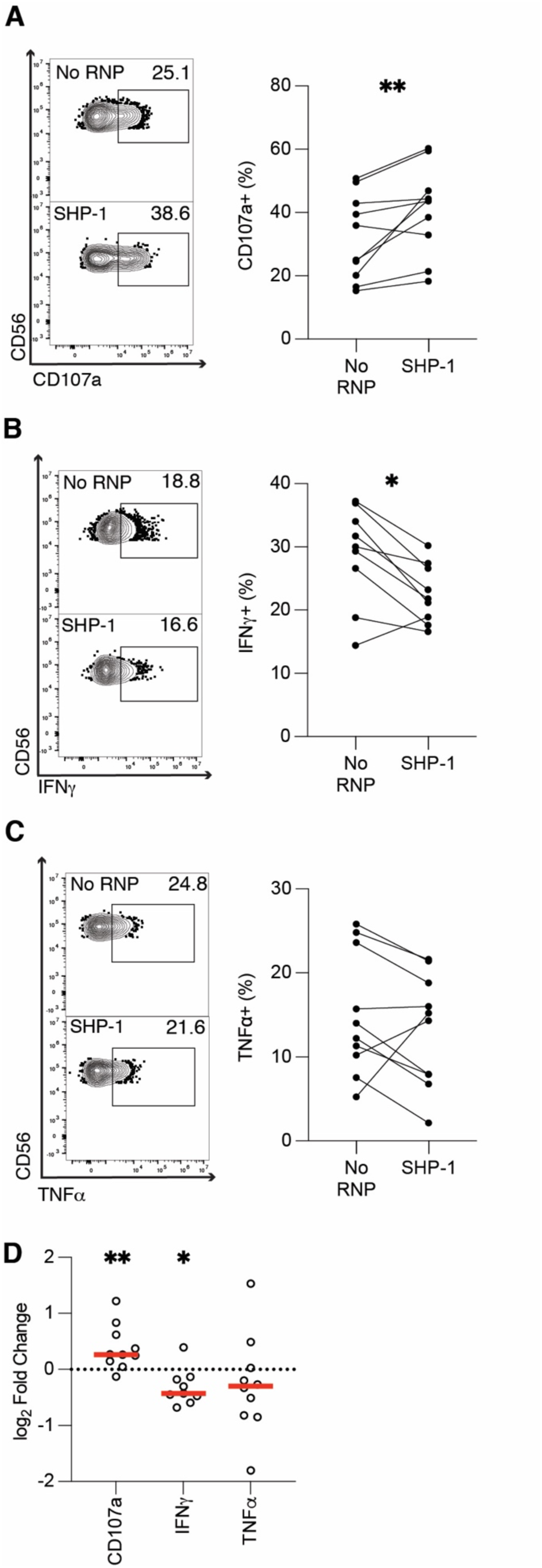
SHP-1 ablation enhances degranulation but diminishes cytokine production. Six days post-CRISPR/Cas9, anti-RBC ADCC assays were run with SHP-1 ablated primary NK cells. CD107a (A), IFNγ (B), and TNFα (C) production from gene-ablated and control primary NK cells was measured by flow cytometry. Flow plots show representative samples for no RNP (top), and SHP-1-ablated (bottom) primary NK cells. Each dot is one subject/donor. Lines connect subjects that were nucleofected without an RNP complex (control) or with an RNP complex. Values for no RNP groups are frequencies from all NK cells, whereas values for the gene-ablated sample are only from cells negative for the protein ablated. CD107a, IFNγ, and TNFα values for each subject shown in A-C were divided by the respective no RNP sample’s value and log-transformed (D). Data from five independent experiments. Wilcoxon matched-pairs signed rank test. *: p < 0.05; **: p < 0.01; lack of stars indicates non-significance.

### SYK and SHP-1 ablation enhance primary NK cell killing

Next, we tested whether our ADCC degranulation results in the anti-RBC assay would correlate with ADCC towards a SKOV-3 target cell line. We hypothesized that gene-ablated cell killing would reflect CD107a expression in the anti-RBC ADCC assays. We used two complementary assays to measure target cell killing— a molecule release ‘DELFIA’ assay and cell tracking with an ‘Incucyte’ machine. In DELFIA assays, SKOV-3 target cells are loaded with a chemical called BADTA, then incubated with effector cells (e.g., NK cells), for 2-6 hours. After incubation, BADTA levels in the supernatant are measured, which reflects pores generated in the target cell or necrotic death releasing the BADTA into the supernatant. For DELFIA assays, we ablated FcRγ/CD3ζ, SYK, ZAP-70, and SHP-1 from primary NK cells. Six days post-CRISPR/Cas9, the gene-ablated cells were incubated with anti-HER2 and BADTA-loaded SKOV-3 cells. In parallel with the anti-RBC results, ablating FcRγ/CD3ζ significantly diminished ADCC function in DELFIA assays (Figure 4A-4D). We found that SYK and SHP-1 ablation resulted in significantly more specific lysis of target cells with 10 ug/mL (Figure 4A-B) and 3 ug/mL anti-HER2 (Figure 4C-D). We also observed significantly less killing in the ZAP-70-ablated samples (4A-4D).

**Figure 4.**
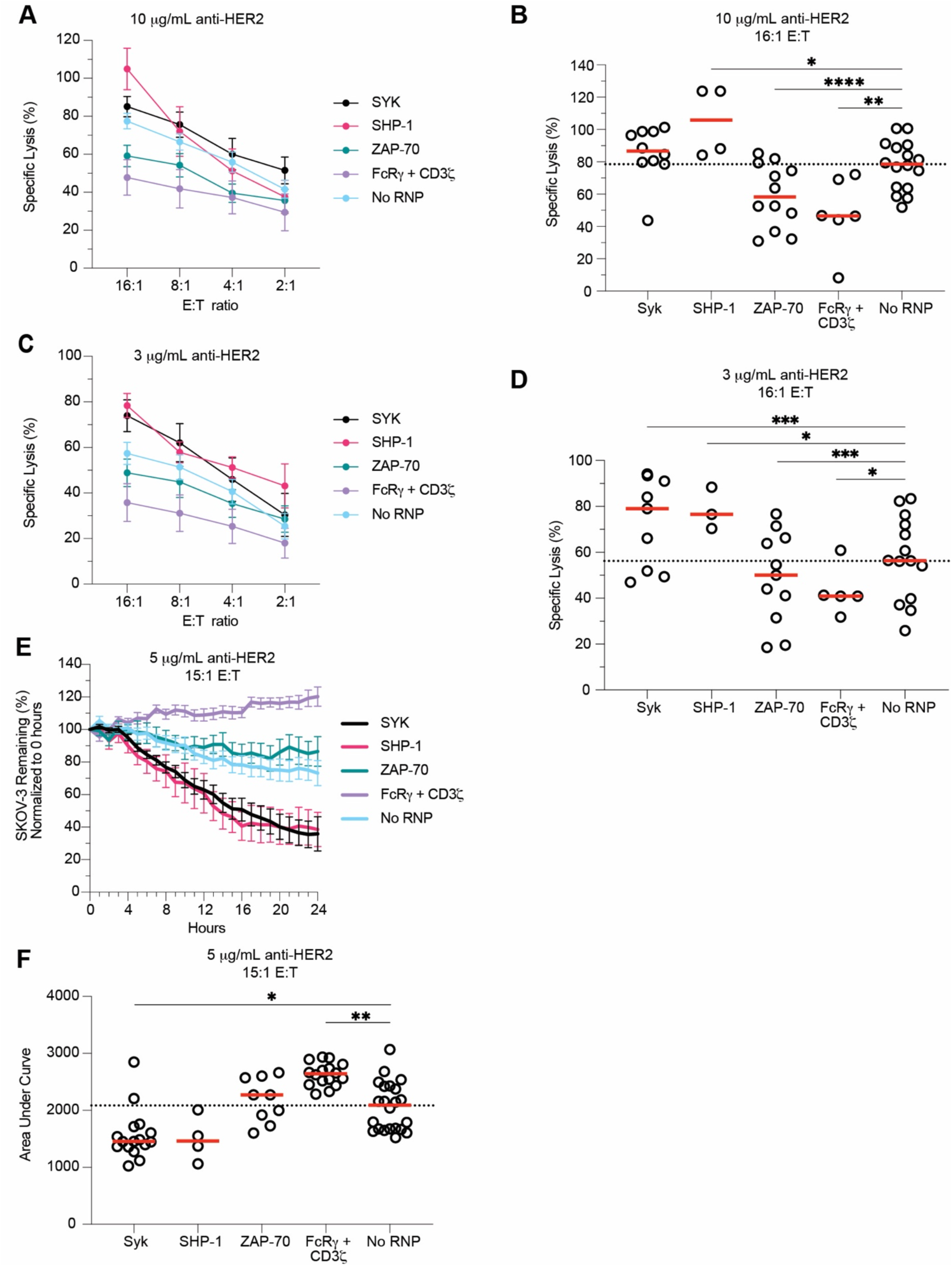
SYK and SHP-1 ablation increases NK cell killing. SYK-, SHP-1-, ZAP-70-, and FcRγ/CD3ζ-ablated primary NK cells were used in DELFIA and Incucyte killing assays six days post-CRISPR/Cas9. DELFIA assays were run by combining NK cells with opsonized SKOV-3 cells. SKOV-3 cells were opsonized with 10 ug/mL (A) and 3 ug/mL (C) anti-HER2, then incubated with NK cells at various E:T ratios. Specific lysis values from both 10 ug/mL (B) and 3 ug/mL (D) samples tested at the 16:1 E:T ratios were plotted. For Incucyte killing assays, NLR-SKOV-3 cells were opsonized with 5 ug/mL anti-HER2 and incubated with primary NK cells at an E:T ratio of about 15:1. Wells were imaged every hour to quantify the percentage of SKOV-3 cells remaining from the original count (E). For each subject, the area under the SKOV-3 % remaining curve was calculated (F). Lines indicate the mean, whereas error bars represent SE (A, C, E). Dots represent individual subjects, whereas red lines represent the median (B, D, F). Data from 7 independent experiments. Dunnet’s multiple comparison’s test for mixed effects model (B, D, F). Numerical values indicate p values for samples with 0.1 > p > 0.05. *: p < 0.05; **: p < 0.01; ***: p < 0.005; ****: p < 0.001. Lack of stars indicates non-significance.

To measure target cell killing over time, we used a microscopy-based live cell tracking Incucyte machine. Fluorescent SKOV-3 cells were opsonized with anti-HER2, incubated with gene-ablated and no RNP NK cells, then imaged every hour for 24 hours. We hypothesized that SKOV-3 death would correlate with degranulation and DELFIA readouts. Once again, we used FcRγ/CD3ζ, SYK, ZAP-70, and SHP-1-ablated cells in Incucyte assays six days post-CRISPR/Cas9. FcRγ/CD3ζ-ablated samples killed significantly fewer target cells (Figure 4E). In line with our other results, we saw that SYK and SHP-1-ablation significantly enhanced killing function (Figure 4E-F). While ZAP-70-ablated samples tended to have a higher median area under the curve (AUC) than control samples, these differences were non-significant (Figure 4E-F). Overall, degranulation, DELFIA specific release, and Incucyte AUC results correlated with each other (Figure S6). Of note, the magnitude of difference between SYK- and SHP-1 ablated cell killing of SKOV-3 cells versus no RNP cells was greater than the difference between the anti-RBC degranulation assay results (Figures 2D, 3D, 4B, D, F). Together, these data support our hypothesis that SYK ablation enhances cytotoxic function in CMV-induced adaptive NK cells.

### SYK-ablated cells retain enhanced function post-expansion

*Ex vivo* expansion of NK cells and subsequent transplantation is an exciting avenue for cancer treatment (Liu et al., 2021). However, the success of these therapies often depends on the ability of NK cells to survive and retain function post-expansion and post-transplant (Miller et al., 2005). Though our main goal was to understand the mechanism by which adaptive NK cells gain enhanced ADCC function, we also wanted to test if SYK-ablated NK cells could survive and retain enhanced function post-expansion. Therefore, we ablated SYK from primary human NK cells, then used them in anti-RBC degranulation and cytokine production assays at days 6, 13, 20, and 27 post-CRISPR. Across nearly all time points, SYK ablation resulted in enhanced CD107a (Figure 5A), IFNγ (Figure 5B), and TNFα (Figure 5C) expression. SYK-ablated samples also retained enhanced killing in DELFIA (Figure 5D-E) and Incucyte (Figure 5F-G) assays with SKOV-3 cells as NK targets. Overall, these results show that SYK ablation’s effect on ADCC function is sustained after expanding the cells for four weeks post-CRISPR, indicating that SYK-negative cells could be considered for NK cell-based ADCC therapies.

**Figure 5.**
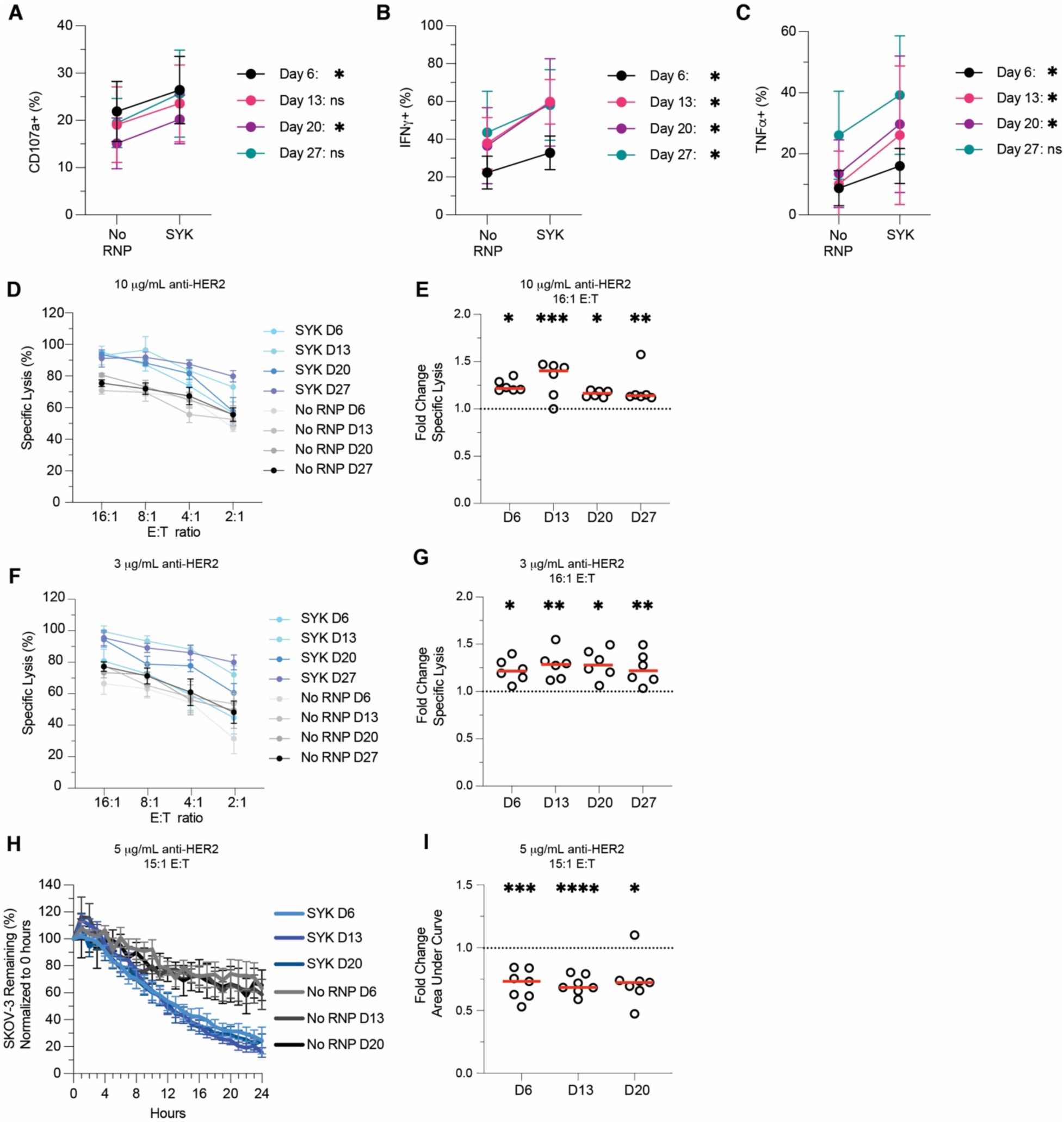
SYK-ablated primary NK cells retain enhanced function. SYK-ablated NK cells were tested in ADCC (A-C), DELFIA (D-G), and Incucyte (H-I) assays at days 6, 13, 20, and 27 post-CRISPR/Cas9. For anti-RBC ADCC assays, CD107a (A), IFNγ (B), and TNFα (C) production in gene-ablated and control primary NK cells was measured by flow cytometry. Plots indicate mean and SD. DELFIA assays were run by incubating NK cells with opsonized SKOV-3 cells. ADCC-specific lysis of SKOV-3 cells opsonized with 10 ug/mL anti-HER2 (D) or 3 ug/mL anti-HER2 (F) at various E:T ratios. SYK-ablated fold change in specific lysis over No RNP group for 16:1 E:T at 10 ug/mL anti-HER2 (E) and 3 ug/mL anti-HER2 (G). Percentage of SKOV-3 cells from t = 0 remaining in Incucyte killing assay (H) and SYK-ablated fold change in SKOV-3 AUC relative to the No RNP group (I). Lines indicate the mean and error bars represent SE (D, F, H). Dots indicate individual subjects and red lines indicate the median (E, G, I). Data from three independent experiments. Wilcoxon matched-pairs signed rank test (A-C). Paired t-tests of raw values (E, G, I). *: p < 0.05; **: p < 0.01; ***: p < 0.005; ****: p < 0.001; lack of stars indicates non-significance.

## Discussion

Adaptive NK cells show enhanced function relative to canonical NK cells (Hwang *et al*., 2012; Lee *et al*., 2015; Schlums *et al*., 2015; Sun *et al*., 2009), and higher frequencies of adaptive NK cells correlate with protection against diseases such as CMV, HIV, and malaria (Cichocki *et al*., 2019; Hart *et al*., 2019; Zhou *et al*., 2015). However, it was unclear which intracellular molecules are required to mediate enhanced adaptive NK cell function and whether the same molecules control both cytotoxicity and cytokine production. Here, we show that ablating SYK kinase in primary NK cells enhances ADCC and cytokine production. SYK-ablated primary NK cells retained their enhanced function up to 27 days post-CRISPR. These results indicate that a mechanism involving SYK kinase—such as ITAM inhibition through SHP-1—could explain why CMV-induced adaptive NK cells show enhanced ADCC and cytokine production relative to conventional NK cells. These results also suggest that SYK-ablated NK cells may have increased ADCC function *in vivo*.

Previous work indicated that adaptive NK cells often lack expression of the transcription factor PLZF and FcRγ. In primary NK cells, loss of FcRγ has been associated with enhanced function, leading to hypotheses that the lack of FcRγ results in a stronger ADCC signaling (Schlums *et al*., 2015; Zhang *et al*., 2013). Additionally, PLZF binds the FcRγ promoter region (Schlums *et al*., 2015). Thus, it seemed possible that loss of PLZF may drive loss of FcRγ, which could lead to enhanced ADCC function. We found that FcRγ ablation led to a modest increase in TNFα production but no significant increase in degranulation or IFNγ expression. In PLZF-ablated samples, neither ADCC nor cytokine production was enhanced for up to 20 days post-CRISPR. Despite these results, PLZF may still be an important transcription factor for adaptive NK cell biology as other transcription factors could have compensated for its loss in our study. In other work, Liu et al. ablated FcRγ and PLZF individually in primary human NK cells and cross-linked CD16A to test ADCC function (Liu *et al*., 2020). Like Liu et al., we observed that ablating PLZF and FcRγ/PLZF did not enhance degranulation or cytokine production during ADCC (Fig. 1). However, Liu et al. also showed that FcRγ-ablation directly enhanced both IFNγ and TNFα production, but not degranulation, during ADCC. By cross-linking CD16A, Liu et al. directly stimulated the CD16A pathway. In contrast to Liu et al., we used two cellular targets to assess ADCC and cytokine production: red blood cells and SKOV-3 cells. Interactions between surface proteins on target and NK cells can modulate NK cell ADCC activity (Lewis et al., 2019; Temming et al., 2019), which may explain the differences between our data and prior work. Given our findings, if gene-ablated NK cells are to be used in clinical settings, this emphasizes the need to test the target cell of interest in pre-clinical assays. Overall, our results show the potential for enhanced cytokine production in FcRγ-negative NK cells but also indicated that lack of the FcRγ chain does not universally enhance ADCC function.

In addition to lacking FcRγ, adaptive NK cells can lack SYK kinase (Lee *et al*., 2015; Schlums *et al*., 2015; Zaghi et al., 2021). Loss of SYK was not observed in adaptive NK cells from malaria subjects in Mali (Hart *et al*., 2019), but has been observed regularly during CMV infection. Like FcRγ-negative cells, SYK-negative primary NK cells exhibit enhanced ADCC function (Lee *et al*., 2015; Schlums *et al*., 2015). In ADCC, SYK preferentially binds the FcRγ chain whereas ZAP-70 preferentially binds the CD3ζ chain (Shiue *et al*., 1995). Thus, it seemed possible that lack of the FcRγ chain in adaptive NK cells results in less SYK-mediated ADCC signaling, indicating a potential role for SYK in controlling ADCC signal strength. SYK ablation enhanced ADCC and cytokine production, whereas ZAP-70 ablation decreased both, indicating that SYK and/or ZAP-70 mediates ADCC signal strength. However, the mechanism behind this finding was initially unclear. The enhanced function of SYK-ablated cells was independent of changes in CD16A expression (Figure S5). This led us to examine SHP-1 regulation of ADCC signal strength as the mechanism. SHP-1 is a phosphatase that can modulate signaling in multiple pathways. Relevant to this study, SHP-1 can dephosphorylate ZAP-70 in T cells (Brockdorff *et al*., 1999; Plas *et al*., 1996). In addition, SHP-1 has been shown to mediate ITAM inhibition (ITAMi), in which SHP-1 is recruited to weakly-ligated Fc receptor ITAMs and dephosphorylates them, dampening signaling (Aloulou *et al*., 2012; Mkaddem *et al*., 2017; Pinheiro da Silva *et al*., 2007). We show that SHP-1 ablation enhances degranulation, indicating that SHP-1 may inhibit ADCC signaling. These results were consistent with enhanced killing by SHP-1-negative NK cells. This implies that ITAMi could be the mechanism behind enhanced ADCC function in SYK and SHP-1-ablated cells. With SYK or SHP-1 ablated, NK cells may signal more extensively through ZAP-70, which—in contrast to SYK—has not been shown to facilitate ITAMi (Ben Mkaddem *et al*., 2014). This could result in the enhanced cytotoxicity we observed. In contrast, with ZAP-70 ablated, SYK may bind FcRγ at a higher rate than with ZAP-70 present. This could facilitate more ITAMi—through SHP-1—than in canonical NK cells, resulting in the decrease in cytotoxic function we observed. This model is depicted in Figure 6. Overall, our results indicate that ITAMi inhibits the ADCC pathway in NK cells, is facilitated by SYK and SHP-1, and could be a mechanism behind the enhanced ADCC observed in adaptive NK cells.

**Figure 6.**
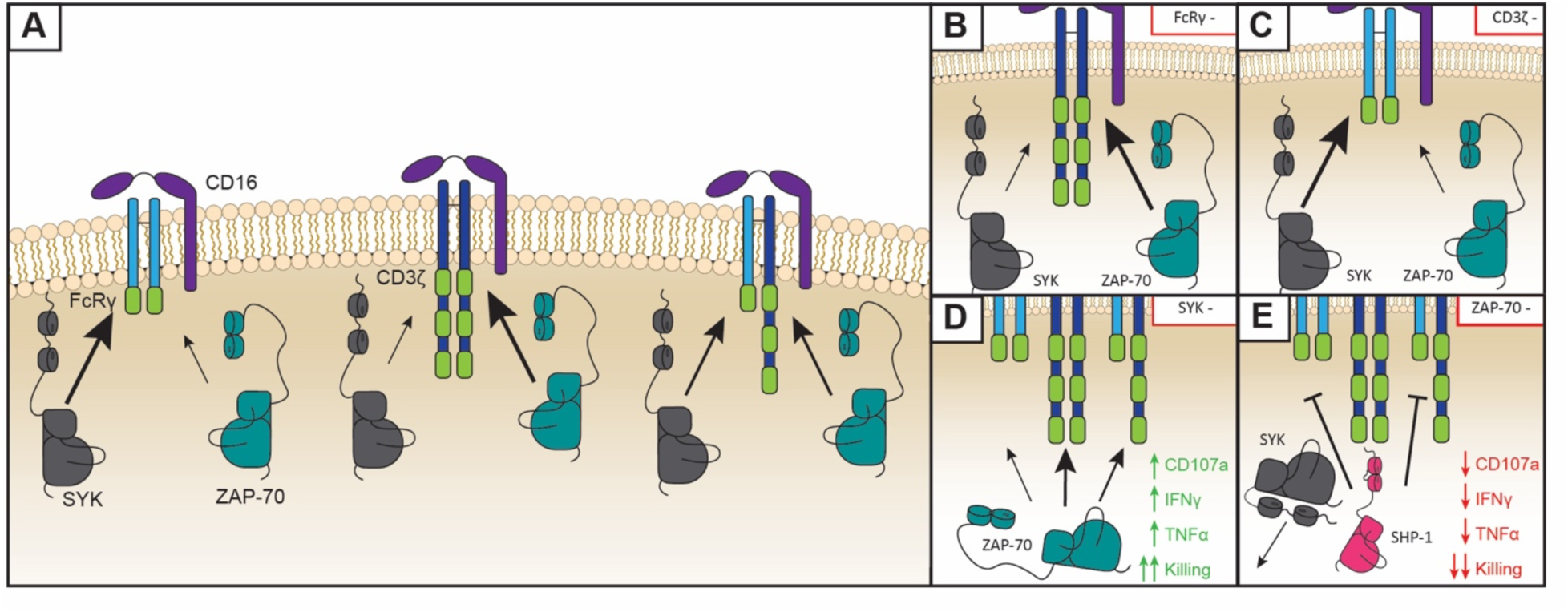
ITAM inhibition model for primary NK cell CD16A signaling. Canonical CD16A signaling pathway (A). In resting NK cells, CD16A constitutively associates with the FcRγ and/or CD3ζ signaling adapters, forming homodimers (left and center) or a heterodimer (right). Crosslinking of CD16A leads to phosphorylation of the ITAM(s) on FcRγ/CD3ζ by Src-family kinases, recruitment of tyrosine kinases SYK/ZAP-70 to the phosphorylated ITAMs, and initiation of downstream cell activation signals. SYK preferentially binds the FcRγ chain (left) whereas ZAP-70 preferentially binds the CD3ζ chain (center). The consequences of signaling chain heterodimer versus homodimer conformations on CD16A signaling remain unclear (right). Effect of FcRγ (B), CD3ζ (C), SYK (D) and ZAP-70 (E) ablation observed on NK cell ADCC and cytokine production. FcRγ-ablation forces NK cells to adopt only CD3ζ homodimer signaling chain conformations (B) whereas CD3ζ-ablation forces FcRγ homodimers (C). We found that neither ablation affects ADCC consistently. SYK-ablation (D) forces ZAP-70- facilitated CD16A signaling, which results in enhanced ADCC, cytokine production, and killing. ZAP-70-ablation (E) results in only SYK binding the signaling adapters. Since SYK has been shown to facilitate ITAM inhibition (ITAMi) via SHP-1, SYK ablation could result in SHP-1 dephosphorylating the signaling chains, resulting in the decreased ADCC, cytokine production, and killing that we observed. This would possibly explain why SYK-ablation (D) results in increased function, as ZAP-70 has not been shown to facilitate ITAMi.

Interestingly, IFNγ production from SHP-1 ablated NK cells was consistently lower than control NK cells—the opposite result of SYK ablation. One explanation could be different CD16A binding thresholds to initiate degranulation versus cytokine production. Fauriat et al. proposed a model for NK cell function in which CD107a signaling requires a low threshold and triggers granule release immediately after stimulation whereas TNFα secretion requires a moderate threshold and longer stimulation (∼3 hours), and IFNγ secretion requires the highest threshold and longest stimulation (∼5.5 hours) (Fauriat et al., 2010). Concurrently, strong activation through CD16A yields a self-regulation loop in which CD16A is enzymatically cleaved by ADAM17 (Jing et al., 2015; Romee *et al*., 2013; Wang *et al*., 2013; Wu et al., 2019). Thus, it’s possible that without SHP-1 negatively regulating CD16A signaling, SHP-1-ablated cells are rapidly activated and degranulate. This could lead to ADAM17 cleaving CD16A from the cell surface, resulting in a lack of signal for eventual IFNγ production. Lack of SHP-1 has been associated with overactive immune cells (Mahmood et al., 2012). Thus, we hypothesize that SHP-1 ablation eliminates inhibition of both SYK (through ITAMi) and ZAP-70 (by dephosphorylation) activity, resulting in robust CD16A signaling, rapid ADAM17-induced cleavage of CD16A, and loss of the hours-long CD16A-stimulation required for IFNγ production. Additional studies beyond the scope of this study are needed to test this hypothesis.

Our results reveal a potential mechanistic explanation for enhanced function in CMV-induced adaptive NK cells. However, they do not explain adaptive NK cell behavior in malaria subjects. Malian adaptive NK cells lack expression of FcRγ and PLZF but were 90% SYK-positive (Hart *et al*., 2019). Like adaptive NK cells in CMV patients, FcRγ-negative cells from malaria subjects showed enhanced ADCC and cytokine production *in vitro* (Hart *et al*., 2019). This highlights the heterogeneity of adaptive NK cells induced by different infections. While our study shows how SYK ablation can enhance degranulation and cytokine production with a CMV-induced adaptive NK cell phenotype, there may be multiple mechanisms driving this enhanced function. Understanding other mechanisms that increase cytotoxicity and cytokine production will be important toward understanding NK cell function and improving their use as a therapeutic intervention.

The CRISPR/Cas9 method we developed allowed us to ablate molecules in the CD16A pathway and define the consequences of the ablations on CD16A signaling in primary human NK cells. These findings build on previous work in the NK and Fc receptor signaling fields as well as adaptive NK cell biology. With gene-editing in primary human NK cells, the field can address new questions on how adaptive NK cells function. Future studies will focus on understanding why SYK ablation enhances NK cell function, ZAP-70 ablation decreases function, and SHP-1 ablation does both. In summary, these findings help define what drives enhanced cytotoxicity and cytokine production in CMV-induced adaptive NK cells and show that SYK ablation may be a viable therapy for CAR NK cells that utilize the CD16A signaling.

## Limitations

Limitations of the study are mentioned throughout the discussion.

## Acknowledgements

This work was funded by the National Institutes of Health (R01 AI146031; R01 AI143828; R21 AI149659) and by the University of Minnesota Clinical and Translational Science Institute (CTSI) and Medical School. Special thanks to Patrick Willey and the rest of the staff at the University of Minnesota University Imaging Centers (UIC, SCR_020997), the University of Minnesota flow cytometry core, and the University of Minnesota Genomics Center. Special thanks to Peter Hinderlie (Dr. Jeffrey Miller’s laboratory) for technical support in setting up Incucyte assays and analyzing results. Thanks to Dr. Peter Morawski (Benaroya Research Institute) for helping with a confirmatory analysis (Infinity Flow R program) for our Biolegend LegendScreen results. Special thanks to the NCI for supplying IL-2 and IL-15 and to Synthego for their guide RNA design tool for CRISPR/Cas9. Thanks to Dr. Jai Rutella and Dr. Miller’s **N**K cell **I**mmunology **R**esearch **F**orum (NIRF) group for their expertise and advice. Thanks to Dr. Joshua Baller for computational and statistical assistance with the data and manuscript. Thanks to Sloane Fussell for providing model figure graphics.

## Author Contributions

Conceptualization J.D.D., J.K.D., J.W., J.M., G.T.H.; Methodology J.D.D., J.K.D., E.P., K.M.S., M.F., J.M., B.Wa., B.We., B.M., G.T.H.; Software J.D.D., C.T.; Validation J.D.D., J.K.D., J.A.S., K.M.S., G.T.H.; Formal Analysis J.D.D., J.K.D.; Investigation J.D.D., J.K.D., J.A.S., J.M.M.; Resources S.E.H., M.F., J.M., B.Wa., B.M., G.T.H.; Data Curation J.D.D., J.K.D., G.T.H.; Writing – Original Draft J.D.D., G.T.H.; Writing – Review & Editing J.D.D., J.K.D., K.M.S., J.W., S.E.H., M.F., J.M., B.Wa., B.We., G.T.H.; Visualization J.D.D., J.K.D.; Supervision S.E.H., B.Wa., B.We., B.M., G.T.H.; Funding Acquisition S.E.H., G.T.H.; Project Administration G.T.H.

## Key resources table

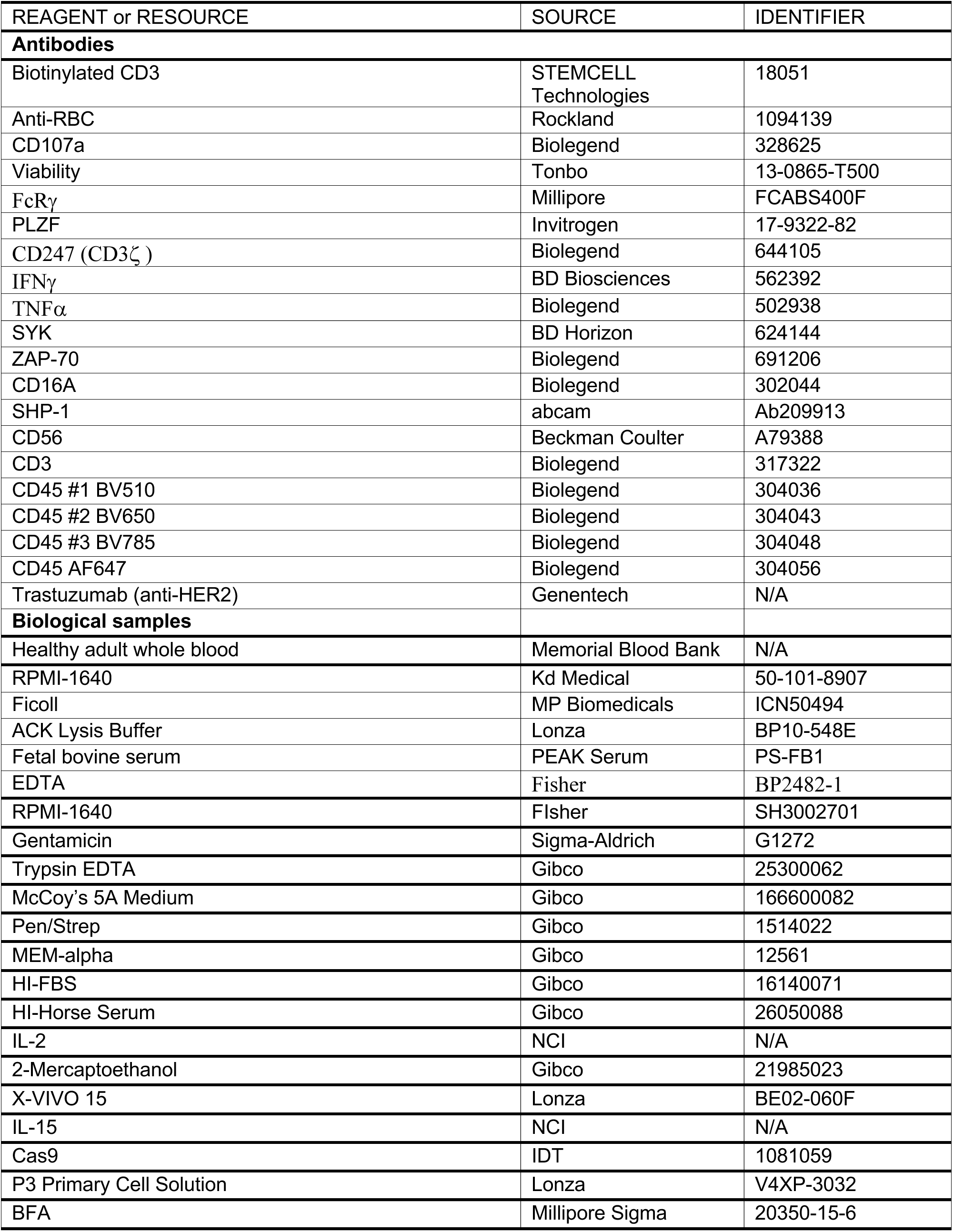
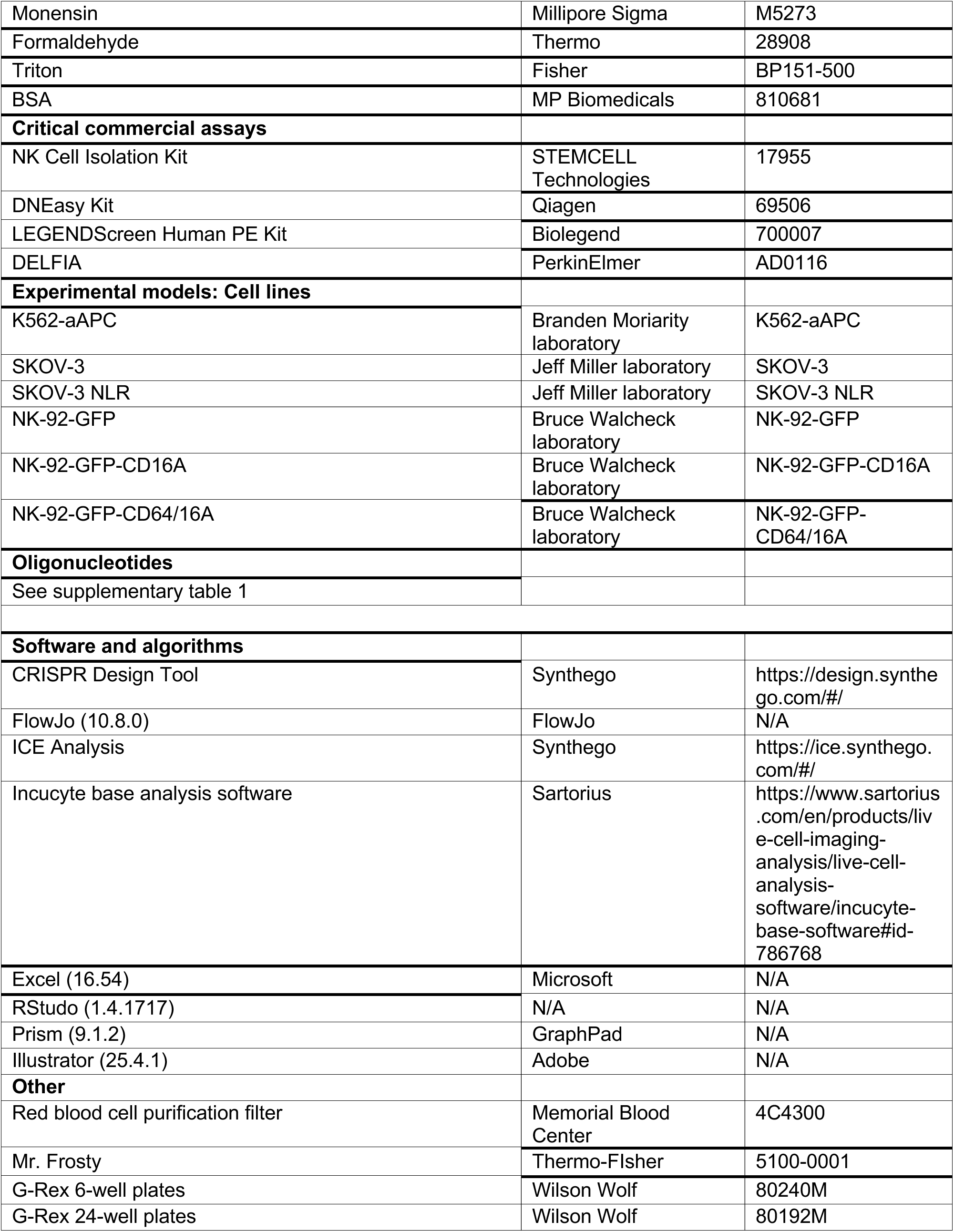

---

**Sup 1.**
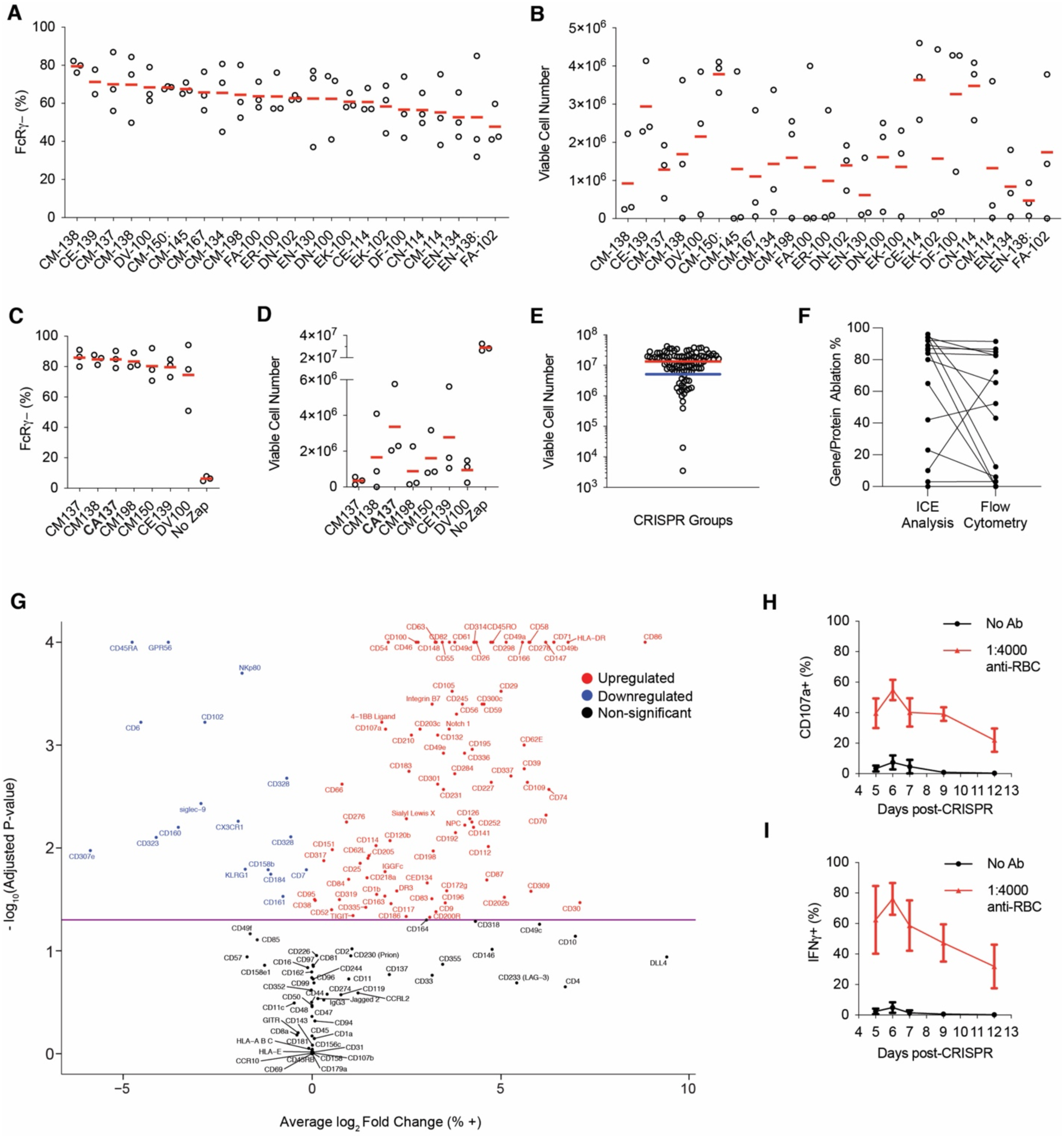
Optimization and validation of the CRISPR/Cas9 gene-editing protocol. Primary NK cell samples (n = 3) were nucleofected with an RNP complex targeting FcRγ. 24 Amaxa-4D Nucleofector codes were used. Six days following CRISPR/Cas9, the samples were stained for FcRγ expression (A) and cell recovery (B). The experiment was repeated with six codes from A-B and CA137, which is in bold (C-D). Each dot is one subject/donor. Red lines represent the median. Primary NK cell samples were nucleofected with a RNP complexes targeting various genes and stained for viability six days post-CRISPR/Cas9 (E). Dots represent individual subject + RNP combinations. The blue line shows the number of cells nucleofected (5×10^6^), whereas the red line shows the average viable cell count six days post-CRISPR/Cas9 (10.7×10^6^). Data from three independent experiments. DNA was isolated from a subset of FcRγ/PLZF-ablated primary NK cell samples six days post-CRISPR/Cas9 and sequenced for ICE analysis alongside staining for FcRγ and PLZF expression (F). ICE analysis values indicate the gene ablation %, whereas flow cytometry values represent the protein ablation % for a sample. Each dot is one subject/donor. Lines connect samples from the same donor + RNP combination. Data from three independent experiments. No RNP cells from days 0 and 7 in the CRISPR/Cas9 protocol were thawed and phenotyped using Biolegend’s LEGENDScreen PE Kit (n = 4, G). Volcano plots show change in expression of all markers from day 0 to day 7. Red dots indicate markers that were significantly upregulated, black dots indicate non-significant changes in expression, and blue dots indicate proteins that were downregulated. Data from one experiment. Two-way ANOVA with Tukey’s multiple comparisons test. No RNP NK cell samples (n = 3) were nucleofected with CA137 and used in anti-RBC ADCC assays at days 5, 6, 7, 9, and 12 post-CRISPR/Cas9. CD107a (H) and IFNγ (I) expression was measured by flow cytometry. Data shown is the average +/- SD of the three subjects at each timepoint for the ADCC (red line) and negative control (black line) groups. Data from one experiment.

**Sup 2.**
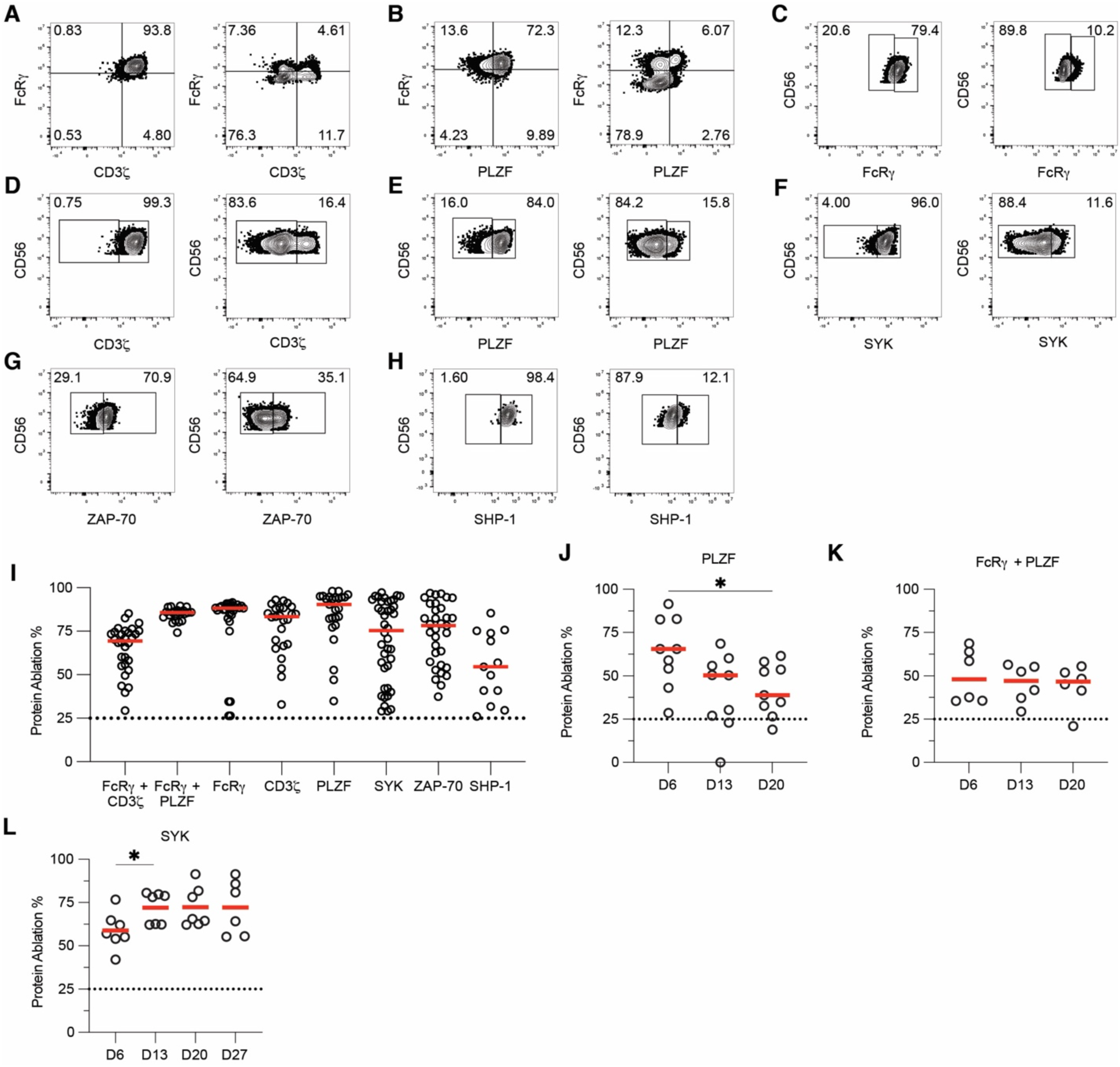
Example flow data and protein ablation for samples. Primary NK cell samples were nucleofected with RNP complexes targeting multiple genes and stained for the targeted protein six days post-CRISPR/Cas9. Representative flow data for each gene targeted are shown (A-H). Protein ablation % for each CRISPR/Cas9 sample six days post-CRISPR (I). The dotted line indicates the minimum protein ablation % for a sample’s data to be used (25%). Red lines represent the median. Each dot represents a data point used in this study. Protein ablation % for PLZF-ablated (J), FcRγ/PLZF-ablated (K), and SYK-ablated (L) samples over time. The dotted line indicates the minimum protein ablation % at day 6 for a sample to be included (25%). Red lines represent the median. One-way ANOVA with Tukey’s multiple comparisons test. *: p < 0.05; lack of stars indicates non-significance.

**Sup 3.**
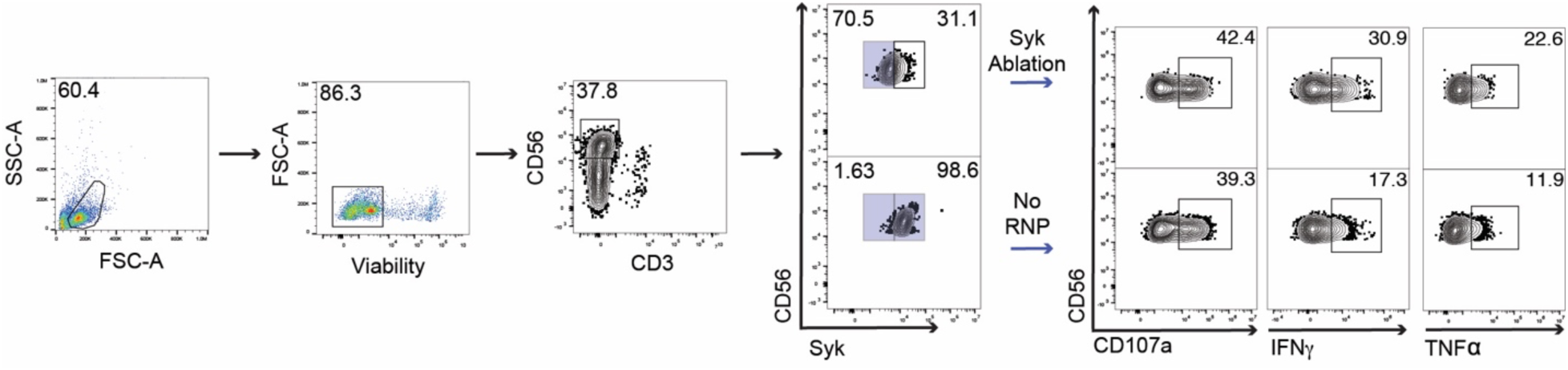
Gating scheme for ADCC assays. Blue boxes around samples indicate the population carried into the next gates.

**Sup 4.**
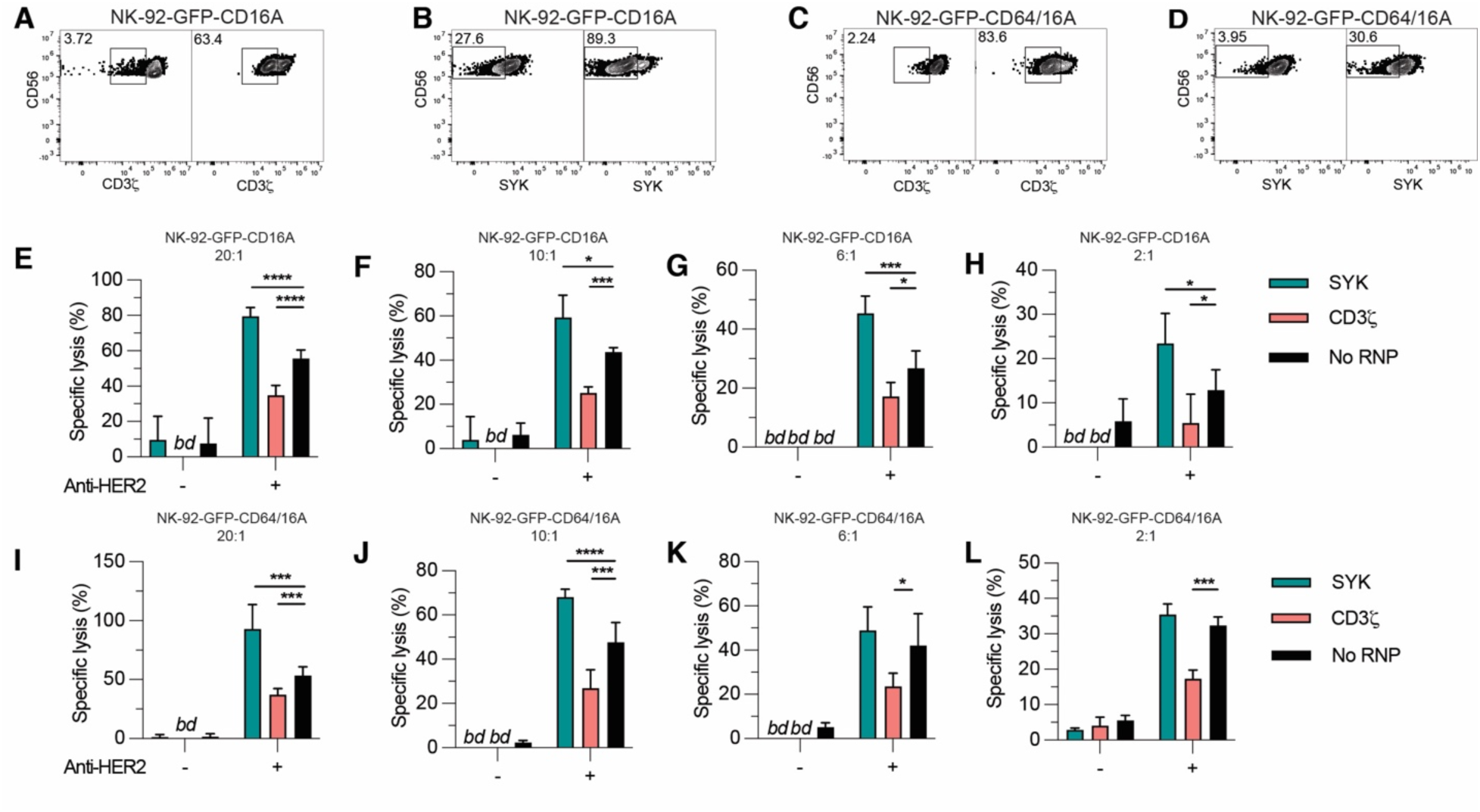
NK-92 ADCC function enhanced by SYK ablation. CD3ζ (A, C) and SYK (Figure B, D) were ablated from NK-92-GFP cells expressing CD16A (A-B, E-H) and CD64/16A (C-D, I-L) via CRISPR/Cas9. Following CRISPR/Cas9, the gene-ablated NK-92 cells were used in DELFIA assays at E:T ratios of 20:1 (E, I), 10:1 (F, J), 6:1 (G, K) and 2:1 (H, L). Technical triplicates were used in the DELFIA assay. Data shown is the specific lysis average + SD in two independent experiments. Values of no lysis are shown as bd (below detection). Paired T-tests between the gene-ablated cell lines and no RNP cell lines. *: p < 0.05; **: p < 0.01; ***: p < 0.005; ****: p < 0.001.

**Sup 5.**
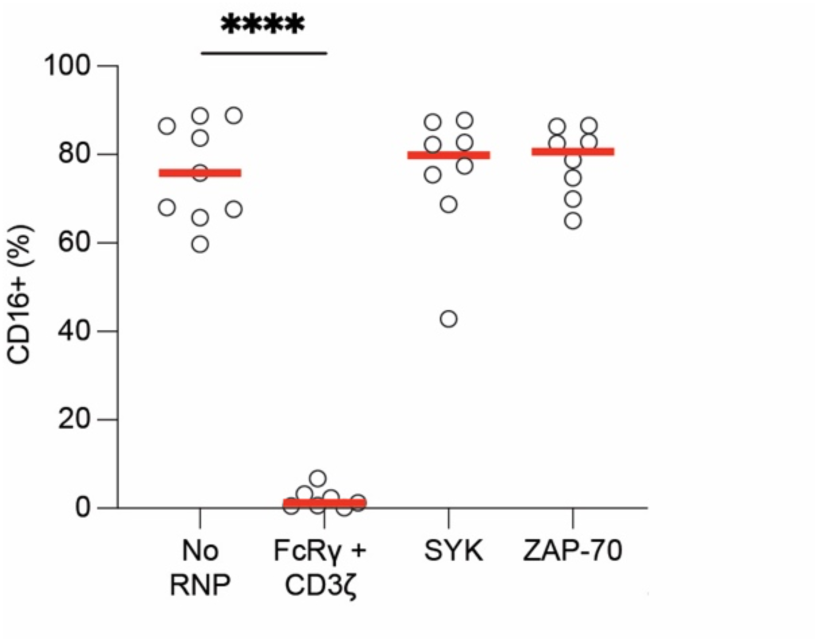
Neither SYK or ZAP-70 ablation affected CD16A surface expression in NK cells. Six days post-CRISPR/Cas9, no RNP and gene-ablated primary NK cell samples were stained for CD16A expression. Dots indicate samples from one subject. Red lines indicate the median. Data shown is from three independent experiments. Dunnet’s multiple comparison’s test for mixed effects model. ****: p < 0.001; lack of stars indicates non-significance.

**Sup 6.**
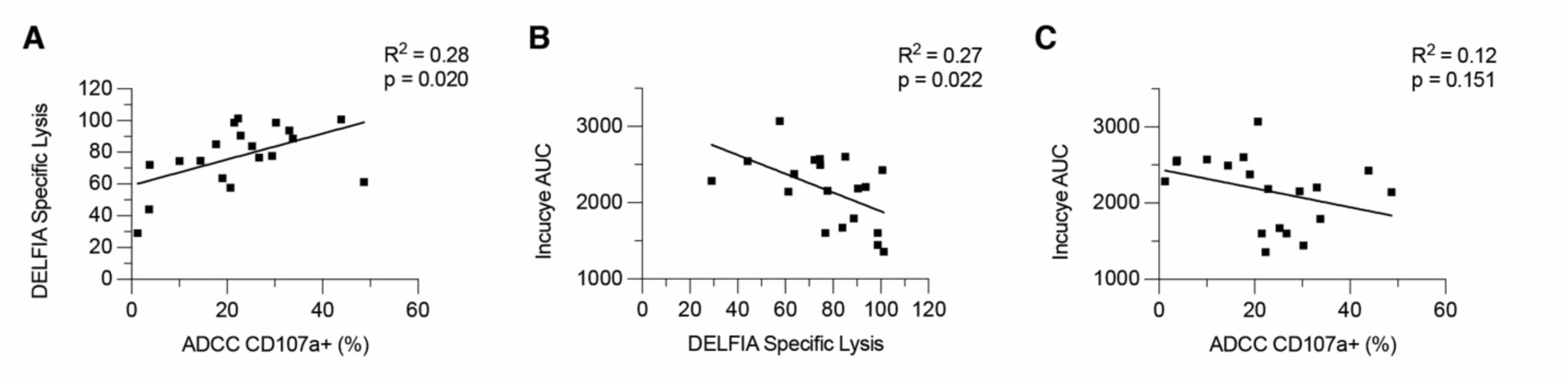
Degranulation, DELFIA, and Incucyte assays correlate. Subject-matched ADCC, DELFIA, and Incucyte assays ran concurrently (n = 14) were used to correlate degranulation/CD107a expression (target cell: RBC), DELFIA (target cell: SKOV-3) specific lysis, and Incucyte (target cell: SKOV-3) area under the curve (AUC) results. Data from four independent experiments. Each dot represents an individual subject. R^2^, p-value, and line of best fit from linear regression analysis shown.

## References

1. Abel, A.M., Yang, C., Thakar, M.S., and Malarkannan, S. (2018). Natural Killer Cells: Development, Maturation, and Clinical Utilization. Front Immunol 9, 1869. 10.3389/fimmu.2018.01869.

2. Abram, C.L., and Lowell, C.A. (2007). The expanding role for ITAM-based signaling pathways in immune cells. Sci STKE 2007, 1–6. 10.1126/stke.3772007re2.

3. Aguilar, O.A., Fong, L.K., Ishiyama, K., DeGrado, W.F., and Lanier, L.L. (2022). The CD3zeta adaptor structure determines functional differences between human and mouse CD16 Fc receptor signaling. J Exp Med 219. 10.1084/jem.20220022.

4. Aloulou, M., Ben Mkaddem, S., Biarnes-Pelicot, M., Boussetta, T., Souchet, H., Rossato, E., Benhamou, M., Crestani, B., Zhu, Z., Blank, U., et al. (2012). IgG1 and IVIg induce inhibitory ITAM signaling through FcgammaRIII controlling inflammatory responses. Blood 119, 3084–3096. 10.1182/blood-2011-08-376046.

5. Arora, G., Hart, G.T., Manzella-Lapeira, J., Doritchamou, J.Y., Narum, D.L., Thomas, L.M., Brzostowski, J., Rajagopalan, S., Doumbo, O.K., Traore, B., et al. (2018). NK cells inhibit Plasmodium falciparum growth in red blood cells via antibody-dependent cellular cytotoxicity. Elife 7. 10.7554/eLife.36806.

6. Ben Mkaddem, S., Hayem, G., Jonsson, F., Rossato, E., Boedec, E., Boussetta, T., El Benna, J., Launay, P., Goujon, J.M., Benhamou, M., et al. (2014). Shifting FcgammaRIIA-ITAM from activation to inhibitory configuration ameliorates arthritis. J Clin Invest 124, 3945–3959. 10.1172/JCI74572.

7. Binyamin, L., Alpaugh, R.K., Hughes, T.L., Lutz, C.T., Campbell, K.S., and Weiner, L.M. (2008). Blocking NK cell inhibitory self-recognition promotes antibody-dependent cellular cytotoxicity in a model of anti-lymphoma therapy. J Immunol 180, 6392–6401. 10.4049/jimmunol.180.9.6392.

8. Bonnin, S. (2020). Introduction to R 2021: 19.11 Volcano Plots. https://biocorecrg.github.io/CRG_RIntroduction/volcano-plots.html.

9. Brockdorff, J., Williams, S., Couture, C., and Mustelin, T. (1999). Dephosphorylation of ZAP-70 and inhibition of T cell activation by activated SHP1. Eur J Immunol 29, 2539–2550.

10. Burrack, K.S., Hart, G.T., and Hamilton, S.E. (2019). Contributions of natural killer cells to the immune response against Plasmodium. Malar J 18, 321. 10.1186/s12936-019-2953-1.

11. Cichocki, F., Taras, E., Chiuppesi, F., Wagner, J.E., Blazar, B.R., Brunstein, C., Luo, X., Diamond, D.J., Cooley, S., Weisdorf, D.J., and Miller, J.S. (2019). Adaptive NK cell reconstitution is associated with better clinical outcomes. JCI Insight 4. 10.1172/jci.insight.125553.

12. Correia, M.P., Stojanovic, A., Bauer, K., Juraeva, D., Tykocinski, L.O., Lorenz, H.M., Brors, B., and Cerwenka, A. (2018). Distinct human circulating NKp30(+)FcepsilonRIgamma(+)CD8(+) T cell population exhibiting high natural killer-like antitumor potential. Proc Natl Acad Sci U S A 115, E5980–E5989. 10.1073/pnas.1720564115.

13. Fauriat, C., Long, E.O., Ljunggren, H.G., and Bryceson, Y.T. (2010). Regulation of human NK-cell cytokine and chemokine production by target cell recognition. Blood 115, 2167–2176. 10.1182/blood-2009-08-238469.

14. Foley, B., Cooley, S., Verneris, M.R., Pitt, M., Curtsinger, J., Luo, X., Lopez-Verges, S., Lanier, L.L., Weisdorf, D., and Miller, J.S. (2012). Cytomegalovirus reactivation after allogeneic transplantation promotes a lasting increase in educated NKG2C+ natural killer cells with potent function. Blood 119, 2665–2674. 10.1182/blood-2011-10-386995.

15. Freedman, T.S., Tan, Y.X., Skrzypczynska, K.M., Manz, B.N., Sjaastad, F.V., Goodridge, H.S., Lowell, C.A., and Weiss, A. (2015). LynA regulates an inflammation-sensitive signaling checkpoint in macrophages. Elife 4. 10.7554/eLife.09183.

16. Goodier, M.R., Lusa, C., Sherratt, S., Rodriguez-Galan, A., Behrens, R., and Riley, E.M. (2016). Sustained Immune Complex-Mediated Reduction in CD16 Expression after Vaccination Regulates NK Cell Function. Front Immunol 7, 384. 10.3389/fimmu.2016.00384.

17. Guma, M., Angulo, A., Vilches, C., Gomez-Lozano, N., Malats, N., and Lopez-Botet, M. (2004). Imprint of human cytomegalovirus infection on the NK cell receptor repertoire. Blood 104, 3664–3671. 10.1182/blood-2004-05-2058.

18. Guma, M., Budt, M., Saez, A., Brckalo, T., Hengel, H., Angulo, A., and Lopez-Botet, M. (2006). Expansion of CD94/NKG2C+ NK cells in response to human cytomegalovirus-infected fibroblasts. Blood 107, 3624–3631. 10.1182/blood-2005-09-3682.

19. Hart, G.T., Tran, T.M., Theorell, J., Schlums, H., Arora, G., Rajagopalan, S., Sangala, A.D.J., Welsh, K.J., Traore, B., Pierce, S.K., et al. (2019). Adaptive NK cells in people exposed to Plasmodium falciparum correlate with protection from malaria. J Exp Med 216, 1280–1290. 10.1084/jem.20181681.

20. Hibbs, M.L., Selvaraj, P., Carpen, O., Springer, T.A., Kuster, H., Jouvin, M.-H.E., and Kinet, J.-P. (1989). Mechanisms for Regulating Expression of Membrane Isoforms of FcyRIII (CD16). Science 246, 1608–1611.

21. Hintz, H.M., Snyder, K.M., Wu, J., Hullsiek, R., Dahlvang, J.D., Hart, G.T., Walcheck, B., and LeBeau, A.M. (2021). Simultaneous Engagement of Tumor and Stroma Targeting Antibodies by Engineered NK-92 Cells Expressing CD64 Controls Prostate Cancer Growth. Cancer Immunol Res 9, 1270–1282. 10.1158/2326-6066.CIR-21-0178.

22. Huang, R.S., Lai, M.C., Shih, H.A., and Lin, S. (2021). A robust platform for expansion and genome editing of primary human natural killer cells. J Exp Med 218. 10.1084/jem.20201529.

23. Hwang, I., Zhang, T., Scott, J.M., Kim, A.R., Lee, T., Kakarla, T., Kim, A., Sunwoo, J.B., and Kim, S. (2012). Identification of human NK cells that are deficient for signaling adaptor FcRgamma and specialized for antibody-dependent immune functions. Int Immunol 24, 793–802. 10.1093/intimm/dxs080.

24. Ivashkiv, L.B. (2011). How ITAMs inhibit signaling. Sci Signal 4, 1–3. 10.1126/scisignal.2001917.

25. Jing, Y., Ni, Z., Wu, J., Higgins, L., Markowski, T.W., Kaufman, D.S., and Walcheck, B. (2015). Identification of an ADAM17 cleavage region in human CD16 (FcgammaRIII) and the engineering of a non-cleavable version of the receptor in NK cells. PLoS One 10, e0121788. 10.1371/journal.pone.0121788.

26. Kanamaru, Y., Pfirsch, S., Aloulou, M., Vrtovsnik, F., Essig, M., Loirat, C., Deschenes, G., Guerin-Marchand, C., Blank, U., and Monteiro, R.C. (2008). Inhibitory ITAM Signaling by FcαRI-FcRγ Chain Controls Multiple Activating Responses and Prevents Renal Inflammation. The Journal of Immunology 180, 2669–2678.

27. Kurosaki, T., and Ravetch, J.V. (1989). A single amino acid in the glycosyl phosphatidylinositol attachment domain determines the membrane topology of Fc gamma RIII. Nature 342, 805–807.

28. Lanier, L.L. (2008). Up on the tightrope: natural killer cell activation and inhibition. Nat Immunol 9, 495–502. 10.1038/ni1581.

29. Lanier, L.L., Yu, G., and Phillips, J.H. (1989). Co-association of CD3z with a receptor (CD16) for IgG Fc on human natural killer cells Nature 342, 803–805.

30. Lanier, L.L., Yu, G., and Phillips, J.H. (1991). ANALYSIS OF FcyRIII (CD16) MEMBRANE EXPRESSION AND ASSOCIATION WITH CD3z AND FceRI-g BY SITE-DIRECTED MUTATION Journal of Immunology 146, 1571–1576.

31. Lee, J., Zhang, T., Hwang, I., Kim, A., Nitschke, L., Kim, M., Scott, J.M., Kamimura, Y., Lanier, L.L., and Kim, S. (2015). Epigenetic modification and antibody-dependent expansion of memory-like NK cells in human cytomegalovirus-infected individuals. Immunity 42, 431–442. 10.1016/j.immuni.2015.02.013.

32. Lewis, G.K., Ackerman, M.E., Scarlatti, G., Moog, C., Robert-Guroff, M., Kent, S.J., Overbaugh, J., Reeves, R.K., Ferrari, G., and Thyagarajan, B. (2019). Knowns and Unknowns of Assaying Antibody-Dependent Cell-Mediated Cytotoxicity Against HIV-1. Front Immunol 10, 1025. 10.3389/fimmu.2019.01025.

33. Liu, S., Galat, V., Galat, Y., Lee, Y.K.A., Wainwright, D., and Wu, J. (2021). NK cell-based cancer immunotherapy: from basic biology to clinical development. J Hematol Oncol 14, 7. 10.1186/s13045-020-01014-w.

34. Liu, W., Scott, J.M., Langguth, E., Chang, H., Park, P.H., and Kim, S. (2020). FcRgamma Gene Editing Reprograms Conventional NK Cells to Display Key Features of Adaptive Human NK Cells. iScience 23, 101709. 10.1016/j.isci.2020.101709.

35. Lopez-Verges, S., Milush, J.M., Schwartz, B.S., Pando, M.J., Jarjoura, J., York, V.A., Houchins, J.P., Miller, S., Kang, S.M., Norris, P.J., et al. (2011). Expansion of a unique CD57(+)NKG2Chi natural killer cell subset during acute human cytomegalovirus infection. Proc Natl Acad Sci U S A 108, 14725–14732. 10.1073/pnas.1110900108.

36. Love, P.E., and Hayes, S.M. (2010). ITAM-mediated signaling by the T-cell antigen receptor. Cold Spring Harb Perspect Biol 2, a002485. 10.1101/cshperspect.a002485.

37. Mahmood, S., Kanwar, N., Tran, J., Zhang, M.L., and Kung, S.K. (2012). SHP-1 phosphatase is a critical regulator in preventing natural killer cell self-killing. PLoS One 7, e44244. 10.1371/journal.pone.0044244.

38. Miller, J.S., Soignier, Y., Panoskaltsis-Mortari, A., McNearney, S.A., Yun, G.H., Fautsch, S.K., McKenna, D., Le, C., Defor, T.E., Burns, L.J., et al. (2005). Successful adoptive transfer and in vivo expansion of human haploidentical NK cells in patients with cancer. Blood 105, 3051–3057. 10.1182/blood-2004-07-2974.

39. Mkaddem, S.B., Murua, A., Flament, H., Titeca-Beauport, D., Bounaix, C., Danelli, L., Launay, P., Benhamou, M., Blank, U., Daugas, E., et al. (2017). Lyn and Fyn function as molecular switches that control immunoreceptors to direct homeostasis or inflammation. Nat Commun 8, 246. 10.1038/s41467-017-00294-0.

40. Monsivais-Urenda, A., Noyola-Cherpitel, D., Hernandez-Salinas, A., Garcia-Sepulveda, C., Romo, N., Baranda, L., Lopez-Botet, M., and Gonzalez-Amaro, R. (2010). Influence of human cytomegalovirus infection on the NK cell receptor repertoire in children. Eur J Immunol 40, 1418–1427. 10.1002/eji.200939898.

41. Mujal, A.M., Delconte, R.B., and Sun, J.C. (2021). Natural Killer Cells: From Innate to Adaptive Features. Annu Rev Immunol 39, 417–447. 10.1146/annurev-immunol-101819-074948.

42. Naeimi Kararoudi, M., Dolatshad, H., Trikha, P., Hussain, S.A., Elmas, E., Foltz, J.A., Moseman, J.E., Thakkar, A., Nakkula, R.J., Lamb, M., et al. (2018). Generation of Knock-out Primary and Expanded Human NK Cells Using Cas9 Ribonucleoproteins. J Vis Exp. 10.3791/58237.

43. Perera Molligoda Arachchige, A.S. (2021). Human NK cells: From development to effector functions. Innate Immun 27, 212–229. 10.1177/17534259211001512.

44. Pinheiro da Silva, F., Aloulou, M., Skurnik, D., Benhamou, M., Andremont, A., Velasco, I.T., Chiamolera, M., Verbeek, J.S., Launay, P., and Monteiro, R.C. (2007). CD16 promotes Escherichia coli sepsis through an FcR gamma inhibitory pathway that prevents phagocytosis and facilitates inflammation. Nat Med 13, 1368–1374. 10.1038/nm1665.

45. Plas, D.R., Johnson, R., Pingel, J.T., Matthews, R.J., Dalton, M., Roy, G., Chan, A.C., and Thomas, M.L. (1996). Direct regulation of ZAP-70 by SHP-1 in T cell antigen receptor signaling. Science 272, 1173–1176.

46. Pomeroy, E.J., Hunzeker, J.T., Kluesner, M.G., Lahr, W.S., Smeester, B.A., Crosby, M.R., Lonetree, C.L., Yamamoto, K., Bendzick, L., Miller, J.S., et al. (2020). A Genetically Engineered Primary Human Natural Killer Cell Platform for Cancer Immunotherapy. Mol Ther 28, 52–63. 10.1016/j.ymthe.2019.10.009.

47. R (2021). R: A language environment for statistical computing. (R Foundation for Statistical Computing, Vienna, Austria).

48. Rautela, J., Surgenor, E., and Huntington, N.D. (2018). Efficient genome editing of human natural killer cells by CRISPR RNP. bioRxiv. 10.1101/406934.

49. Romee, R., Foley, B., Lenvik, T., Wang, Y., Zhang, B., Ankarlo, D., Luo, X., Cooley, S., Verneris, M., Walcheck, B., and Miller, J. (2013). NK cell CD16 surface expression and function is regulated by a disintegrin and metalloprotease-17 (ADAM17). Blood 121, 3599–3608. 10.1182/blood-2012-04-425397.

50. Schlums, H., Cichocki, F., Tesi, B., Theorell, J., Beziat, V., Holmes, T.D., Han, H., Chiang, S.C., Foley, B., Mattsson, K., et al. (2015). Cytomegalovirus infection drives adaptive epigenetic diversification of NK cells with altered signaling and effector function. Immunity 42, 443–456. 10.1016/j.immuni.2015.02.008.

51. Shiue, L., Green, J., Green, O.M., Karas, J.L., Morgenstern, J.P., Ram, M.K., Taylor, M.K., Zoller, M.J., Zydowsky, L.D., Bolen, J.B., and Brugge, J.S. (1995). Interaction of p72syk with the gamma and beta Subunits of the High-Affinity Receptor for Immunoglobulin E, FcεRI. Molecular and Cellular Biology 15, 272–281.

52. Snyder, K.M., Hullsiek, R., Mishra, H.K., Mendez, D.C., Li, Y., Rogich, A., Kaufman, D.S., Wu, J., and Walcheck, B. (2018). Expression of a Recombinant High Affinity IgG Fc Receptor by Engineered NK Cells as a Docking Platform for Therapeutic mAbs to Target Cancer Cells. Front Immunol 9, 2873. 10.3389/fimmu.2018.02873.

53. Sun, J.C., Beilke, J.N., and Lanier, L.L. (2009). Adaptive immune features of natural killer cells. Nature 457, 557–561. 10.1038/nature07665.

54. Temming, A.R., de Taeye, S.W., de Graaf, E.L., de Neef, L.A., Dekkers, G., Bruggeman, C.W., Koers, J., Ligthart, P., Nagelkerke, S.Q., Zimring, J.C., et al. (2019). Functional Attributes of Antibodies, Effector Cells, and Target Cells Affecting NK Cell-Mediated Antibody-Dependent Cellular Cytotoxicity. J Immunol 203, 3126–3135. 10.4049/jimmunol.1900985.

55. Wang, Y., Wu, J., Newton, R., Bahaie, N.S., Long, C., and Walcheck, B. (2013). ADAM17 cleaves CD16b (FcgammaRIIIb) in human neutrophils. Biochim Biophys Acta 1833, 680–685. 10.1016/j.bbamcr.2012.11.027.

56. Watzl, C., and Long, E.O. (2010). Signal transduction during activation and inhibition of natural killer cells. Curr Protoc Immunol Chapter 11, Unit 11 19B. 10.1002/0471142735.im1109bs90.

57. Wu, J., Mishra, H.K., and Walcheck, B. (2019). Role of ADAM17 as a regulatory checkpoint of CD16A in NK cells and as a potential target for cancer immunotherapy. J Leukoc Biol 105, 1297–1303. 10.1002/JLB.2MR1218-501R.

58. Yang, C., Siebert, J.R., Burns, R., Gerbec, Z.J., Bonacci, B., Rymaszewski, A., Rau, M., Riese, M.J., Rao, S., Carlson, K.S., et al. (2019). Heterogeneity of human bone marrow and blood natural killer cells defined by single-cell transcriptome. Nat Commun 10, 3931. 10.1038/s41467-019-11947-7.

59. Zaghi, E., Calvi, M., Puccio, S., Spata, G., Terzoli, S., Peano, C., Roberto, A., De Paoli, F., van Beek, J.J., Mariotti, J., et al. (2021). Single-cell profiling identifies impaired adaptive NK cells expanded after HCMV reactivation in haploidentical HSCT. JCI Insight 6. 10.1172/jci.insight.146973.

60. Zhang, T., Scott, J.M., Hwang, I., and Kim, S. (2013). Cutting edge: antibody-dependent memory-like NK cells distinguished by FcRgamma deficiency. J Immunol 190, 1402–1406. 10.4049/jimmunol.1203034.

61. Zhou, J., Amran, F.S., Kramski, M., Angelovich, T.A., Elliott, J., Hearps, A.C., Price, P., and Jaworowski, A. (2015). An NK Cell Population Lacking FcRgamma Is Expanded in Chronically Infected HIV Patients. J Immunol 194, 4688–4697. 10.4049/jimmunol.1402448.

